# Defining the Subcellular Distribution and Metabolic Channeling of Phosphatidylinositol

**DOI:** 10.1101/677229

**Authors:** Joshua G. Pemberton, Yeun Ju Kim, Nivedita Sengupta, Andrea Eisenreichova, Daniel J. Toth, Evzen Boura, Tamas Balla

**Affiliations:** Section on Molecular Signal Transduction, Eunice Kennedy Shriver National Institute of Child Health and Human Development, National Institutes of Health, Bethesda, MD, USA; Institute of Organic Chemistry and Biochemistry, Czech Academy of Sciences, Prague, Czech Republic

**Keywords:** phosphatidylinositol, polyphosphoinositides, membrane biology, phospholipid metabolism, lipid transport, signal transduction.

## Abstract

Phosphatidylinositol (PtdIns) is an essential structural component of eukaryotic membranes that also serves as the common precursor for polyphosphoinositide (PPIn) lipids. Despite the recognized importance of PPIn species for signal transduction and membrane homeostasis, there is still a limited understanding of how the dynamic regulation of PtdIns synthesis and transport contributes to the turnover of PPIn pools. To address these shortcomings, we capitalized on the substrate selectivity of a bacterial enzyme, PtdIns-specific PLC, to establish a molecular toolbox for investigations of PtdIns distribution and availability within intact cells. In addition to its presence within the ER, our results reveal low steady-state levels of PtdIns within the plasma membrane (PM) and endosomes as well as a relative enrichment of PtdIns within the cytosolic leaflets of the Golgi complex, peroxisomes, and outer mitochondrial membranes. Kinetic studies also demonstrate the requirement for sustained PtdIns supply from the ER for the maintenance of monophosphorylated PPIn species within the PM, Golgi complex, and endosomal compartments.

**Summary:** Pemberton et al. characterize a molecular toolbox for the visualization and manipulation of phosphatidylinositol (PtdIns) within intact cells. Results using these approaches define the steady-state distribution of PtdIns across subcellular membrane compartments as well as provide new insights into the relationship between PtdIns availability and polyphosphoinositide turnover.

## Introduction

The dynamic remodeling of cellular membranes relies on molecular mechanisms that exploit the unique physiochemical properties of individual lipid species (van Meer et al., 2008; Holthuis and Menon, 2010). Additionally, the local lipid composition, in concert with integral and peripheral proteins, effectively defines membrane characteristics such as fluidity, thickness, and surface charge (Bigay and Antonny, 2012; Drin, 2014; Barelli and Antonny, 2016; Jackson et al., 2016). The ability to maintain specific membrane compartments with distinct protein and lipid compositions directly facilitates the spatial organization of metabolic functions that underlie cellular homeostasis. Our current understanding of membrane composition has benefitted greatly from biomolecular separation techniques such as high-performance thin-layer chromatography, as well as, more recently, though the establishment of comprehensive lipidomics methodologies (Shevchenko and Simons, 2010; Wenk, 2010; Harkewicz and Dennis 2011; Tumanov and Kamphorst, 2017). However, these approaches lack spatial information related to how lipid species are distributed or transported throughout the membrane compartments that exist within an intact cell. While subcellular fractionation studies have provided the first detailed information regarding the lipid composition of specific organelles (van Meer and de Kroon, 2011; Klose et al., 2013; Brügger, 2014), it is becoming increasingly evident that even the most efficient isolation methods cannot work quickly enough to preserve the lipid composition of cellular membranes and also suffer from the unavoidable cross-contamination of organelles (van Meer, 2005; Satori et al., 2013; Kappler et al., 2016). Outside of these *in vitro* biochemical studies, imaging breakthroughs using endogenous membrane-binding protein domains have allowed for the visualization of embedded lipid species with high spatial and temporal resolution in live cells (Varnai et al., 2017; Wills et al., 2018). The refinement and rational design of lipid biosensors has been aided by detailed structural and biophysical studies that have established sequence features that directly contribute to the specificity and affinity of peripheral membrane-binding protein domains based on the recognition of general membrane features as well as individual lipid components (Lee, 2003; Cho and Stahelin, 2005; Pogozheva et al., 2013; Whited and Johs, 2015; Pasenkiewicz-Gierula et al., 2016). Despite these progresses, imaging applications are currently limited to only a few classes of lipids and have been predominantly focused on lipids involved in cellular signaling. Visualization or manipulation of structural lipids that form the bulk of eukaryotic membranes is still to be accomplished. In particular, among the core structural lipids, phosphatidylinositol (PtdIns) is unique in that it also serves as the precursor for all polyphosphoinositides (PPIn), which are known to directly control important aspects of membrane trafficking and cellular metabolism (Balla, 2013).

PtdIns synthesis occurs through the conjugation of *myo*-inositol to a cytidine diphosphate (CDP)- activated diacylglycerol (DAG) backbone and is dependent upon the activity of PtdIns synthase (PIS, also called CDIPT; Agranoff et al., 1958; Paulus and Kennedy, 1960; Tanaka et al., 1996). PIS is an integral membrane protein with undisputed activity associated with the ER (Williamson and Moore, 1976; Morris et al., 1990), although previous work from our group has also shown that, in mammalian cells, catalytically-active PIS, but not inactive mutants, is enriched within a mobile ER-derived sub-compartment that may function to actively distribute PtdIns to subcellular membranes (Kim et al., 2011). Within specific membrane compartments, modification of PtdIns is tightly coordinated by substrate-selective kinases and phosphatases, whose integrated activity determines the local phosphorylation status of the *myo*-inositol headgroup at the 3-, 4-, and 5-positions to generate seven distinct PPIn species: including mono-, bis-, and tris-phosphorylated PtdIns derivatives (Balla, 2013; Hsu and Mao, 2015; Burke, 2018). Individual PPIn isomers localize to overlapping as well as distinct membrane compartments where they not only contribute to the assembly and initiation of signaling responses, but also function to maintain membrane identity (Hammond and Balla, 2015; Pemberton and Balla, 2018; Dickson and Hille, 2019). Due to the expansive cellular roles of PPIn lipids, both as substrates for signaling enzymes and sites for protein-membrane interactions, it is not surprising that many human pathologies are caused by perturbations in PPIn production or turnover (Pendaries et al., 2003; Wymann and Schneiter, 2008; McCrea and De Camilli, 2009; Bunney and Katan, 2010; Dyson et al., 2012; Altan-Bonnet, 2017). The interest in PPIn metabolism has resulted in significant efforts to validate selective biosensors to define the distribution and inter-conversion pathways that modulate localized PPIn levels (Indevall-Hagren and De Camilli, 2014; Varnai et al., 2017; Wills et al., 2018). However, as the complex processes governing PPIn metabolism must begin with PtdIns synthesis, investigations of PPIn metabolism continue to be limited by the lack of understanding of how PtdIns production and transport are controlled. This is due in large part to the fact that mammalian proteins with enough specificity and affinity to be used as PtdIns-selective probes have yet to be identified. That said, Gram-positive bacteria produce virulence factors that directly target PtdIns in host membranes, including the ancestral homologs of the PLC superfamily. Unlike the canonical PLCs from higher vertebrates that use PtdIns(4,5)P_2_ as a substrate and require Ca^2+^ as a co-factor, the catalysis carried out by bacterial PtdIns-specific PLC (PI-PLC)s typically occurs independent of metal ions and these enzymes specifically hydrolyze PtdIns to generate lipid-soluble DAG and water-soluble inositol 1-phosphate (Bruzik and Tsai, 1994; Williams and Katan, 1996; Heinz et al., 1998). *In vivo*, most forms of bacterial PI-PLC work extracellularly by cleaving the headgroup from glycosyl-PtdIns (GPI) linkages that anchor GPI-targeted proteins to the outer leaflet of the plasma membrane (PM; Low and Saltiel, 1988; Ferguson and Williams, 1988; Sharom and Lehto, 2002); although intracellular functions of these enzymes have also been documented (Wadsworth and Goldfine, 1999; Wei et al., 2005; Poussin et al., 2009). Consequently, extensive structural and biochemical investigations of PI-PLCs were undertaken in hopes of generating strain-specific antibiotics or small molecules that could selectively target these secreted enzymes (Vinod et al., 1994; Martin and Wagman, 1996; Morris et al., 1996). Moreover, PI-PLCs were also recognized as ideal models for understanding how peripheral membrane proteins interact with membrane surfaces to carryout catalysis (Griffith and Ryan, 1999; Roberts et al., 2018). Collectively, these detailed *in vitro* analyses have provided a comprehensive description of the enzymatic activity and membrane-binding strategies utilized by PI-PLCs. These studies also highlighted the potential suitability of these enzymes as protein scaffolds for the design of PtdIns-selective molecular tools.

In the present study, we chose to revisit the utility of the well-studied and PtdIns-selective bacterial PI-PLCs for designing tools to visualize or manipulate PtdIns content in the membranes of intact cells. Using the structural descriptions available for PI-PLCs, we selected the highly active enzyme from *Bacillus cereus* (Kuppe et al., 1989; Volwerk et al., 1989) to use as a platform for protein engineering. Importantly, the *Bacillus cereus (Bc*)PI-PLC shows remarkable specificity for PtdIns and does not recognize or show catalytic activity towards phosphorylated PPIn species or other common phospholipids (Ikezawa and Taguchi, 1981). In fact, the substrate selectivity of the *Bacillus* PI-PLCs even extends to the physicochemistry of the inositol ring, where the *myo* stereoisomer of inositol is preferentially hydrolyzed and the naturally occurring 1D configuration of the esterified headgroup is an absolute requirement for catalysis (Leigh et al., 1992; Lewis et al., 1993; Bruzik et al., 1994). Just as importantly, variations to the glycerol backbone and fatty acyl chains are well tolerated by the *Bc*PI-PLC (Guther et al., 1994). Capitalizing on these features, our general strategy was two-fold: first, we targeted residues within the conserved catalytic triad that would abolish enzymatic activity, but maintain substrate coordination within the active site in order to mark the intracellular distribution of PtdIns. Second, we generated *Bc*PI-PLC constructs with minimal interfacial binding, and therefore low basal catalytic activity from the cytosol. Such modified enzymes would still be capable of rapidly hydrolyzing PtdIns when recruited in the proximity of membrane-embedded substrate. Our results using catalytic mutants of the *Bc*PI-PLC suggest that PtdIns is localized to ER, the biochemically-defined site of PtdIns synthesis, but is also enriched in the cytosolic leaflets of the Golgi complex, peroxisomes, and mitochondria. Strikingly, we did not observe significant localization of the *Bc*PI-PLC variants within the PM or to endosomal compartments in any of the mammalian cell types examined. The membrane distribution of PtdIns was further investigated using recruitable versions of the modified *Bc*PI-PLC using organelle-targeted anchors and a chemically-inducible protein heterodimerization system. These enzyme recruitment studies show the rapid and localized production of DAG, the direct cleavage product of PtdIns hydrolysis, on the ER, mitochondria, peroxisome, and Golgi complex; as well as, to a lesser extent in Rab5- and Rab7-positive endosomes, but not within the PM. Importantly, this alternative strategy using *Bc*PI-PLC-mediated DAG production as an indirect proxy of PtdIns membrane contents confirmed the PtdIns distributions mapped by the *Bc*PI-PLC-based localization probe. Lastly, we also developed bioluminescence energy-transfer (BRET)-based biosensors to monitor organelle-specific PtdIns, DAG, and PPIn dynamics at the level of cell populations and combined this method with the use of the recruitable *Bc*PI-PLC construct to characterize the role of PtdIns availability and supply for the generation of PPIn species within distinct membrane compartments of live cells. These studies reveal the explicit need for the sustained delivery of PtdIns from the ER, rather than the absolute steady-state content of PtdIns, for the maintenance of monophosphorylated PPIn species within the PM, Golgi complex, and endosomal compartments. Overall, our findings support an important role for PtdIns transfer and substrate channeling in the spatial control of PPIn metabolism.

## Results

### Visualizing the membrane distribution of PtdIns using the PI-PLC scaffold

Unlike the low-abundance PPIn lipids, the intracellular distribution of PtdIns has never been observed in live cells. Here, we devised a strategy to visualize the steady-state distributions of PtdIns that capitalizes on the substrate selectivity and high specific activity of bacterial PI-PLCs. PI-PLCs are small monomeric proteins, roughly 300 amino acid residues in length, that consist of a single polypeptide with no disulfide bonds (Griffith et al., 1991); features that make them ideally suited to use as a template for developing an imaging probe or engineered enzyme. Despite clear effects on cellular DAG and PPIn distributions, our previous studies expressing the PI-PLC from *Listeria monocytogenes* failed to show any membrane localization of the enzyme; presumably because of insufficient binding affinity (Fig.1A; Kim et al., 2011). Therefore, we have turned to another PI-PLC isozyme from *Bacillus cereus* that shows enhanced catalytic activity *in vitro* (Gandhi et al., 1993; Bruzik and Tsai, 1994) and, based on the available crystal structures (Heinz et al., 1995; Heinz et al., 1996; Moser et al., 1997), possesses a more defined pocket for accommodating the inositol head group as well as protruding hydrophobic elaborations present at the membrane-binding interface (Fig.1A; highlighted in yellow). Briefly, the gene sequence for the *Bc*PI-PLC (residues 32-329) was codon-optimized and prepared by custom synthesis within a standard plasmid shuttle before sub-cloning into mammalian expression vectors as a fusion to the C-terminus of EGFP. In all instances, the N-terminal hydrophobic signal sequence (residues 1-31) was removed to prevent targeting of the *Bc*PI-PLCs into the secretory pathway. Expression of the GFP-*Bc*PI-PLC in mammalian cells was extremely cytotoxic, with only small necrotic cells remaining 16-20 h post-transfection (Fig.1A). Mutagenesis studies targeting residues within the active site, including the conserved catalytic triad, were used to inactivate the GFP-*Bc*PI-PLC in hopes of revealing any associations with subcellular membranes. Early studies of *Bc*PI-PLC enzymology identified His32 as the general base that is responsible for abstracting the hydrogen from the C2 hydroxyl on the inositol ring (Fig.1B; Gässler et al., 1997; Hondal et al., 1998). Swapping His32 for alanine (H32A) eliminated the clear cytotoxicity associated with the fully-active *Bc*PI-PLC; suggesting that this mutation effectively prevents *Bc*PI-PLC catalysis (Fig.1C). Importantly, expression of the GFP-*Bc*PI-PLC^H32A^ also revealed a weak association with intracellular membranes, which could reflect the site of enzymatic activity and therefore substrate availability. In an attempt to enhance the relative affinity of this probe, we targeted a second histidine, His82, which also plays a central role in the catalytic mechanism as part of core catalytic triad, but does not directly form hydrogen-bonds with the inositol headgroup (Fig.1B; Gässler et al., 1997; Hondal et al., 1998). Targeting this residue would seem to not only abolish catalytic activity (Gässler et al., 1997), but would also maintain the relative affinity of the active site for PtdIns by preserving all of the residues responsible for coordination of the bound substrate. In line with this idea, relative to the GFP-*Bc*PI-PLC^H32A^, we observed a clear increase in the membrane association of the putative GFP-*Bc*PI-PLC^H82A^ substrate trap, as evidenced from the limited fraction of the signal present within the cytosol (Fig.1D). Importantly, structural analysis of the resulting *Bc*PI-PLC^H82A^ variant using X-ray crystallography showed no alterations to the overall architecture of the active site or any gross changes to either the protein stability or membrane-binding interface (Fig.S1). Detailed co-localization studies using GFP-*Bc*PI-PLC^H82A^ are presented in more detail as part of the subsequent sections, but, generally, we observed specific association of the probe with the ER (Fig.3A), as well as marked enrichments in the mitochondria (Fig.4A), peroxisomes (Fig.5A and 5B), and Golgi complex (Fig.6A and 6B). Strikingly, the GFP-*Bc*PI-PLC^H82A^ probe did not localize to the PM (Fig.8A) or to various endosomal compartments (Fig.10A and 10D). The steady-state membrane distribution of GFP-*Bc*PI-PLC^H82A^ was consistent across representative mammalian cell lines, including the COS-7, HEK293, and HT-1080 lineages (Fig.S2). To further validate these localization studies, we looked to establish a novel heterodimerization system using the active *Bc*PI-PLC for the rapid hydrolysis of PtdIns at precise locations throughout the endomembrane system in live cells.

**Figure 1.**
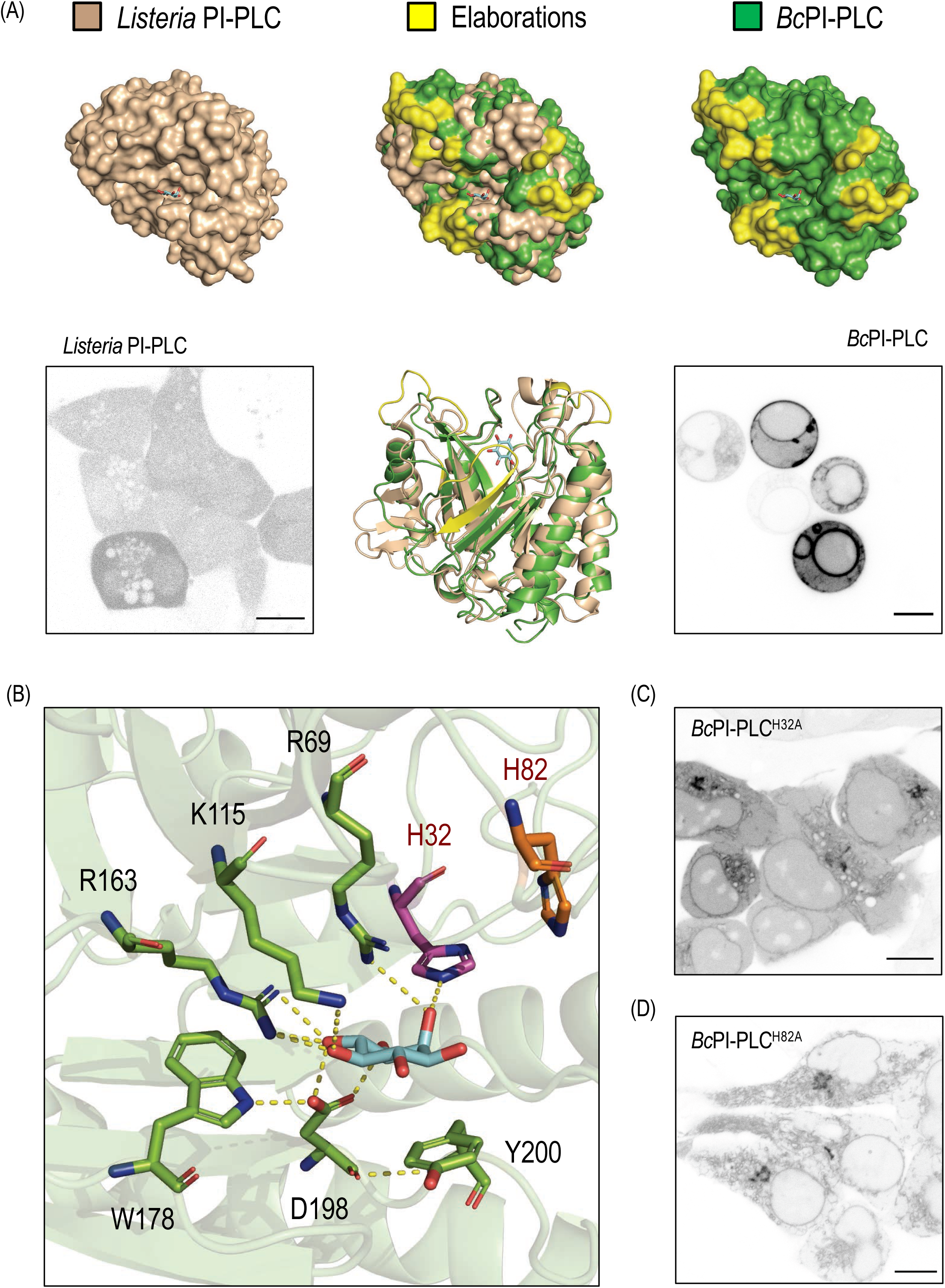
Visualizing the subcellular distribution of PtdIns using the *Bc*PI-PLC scaffold. (A) Structural comparison of the *Listeria monocytogenes* (top row, left panel, beige; PDB Accession: 1AOD; Moser et al., 1997) and *Bacillus cereus* (top row, right panel,green; PDB Accession: 1PTG; Heinz et al., 1995) PI-PLCs bound to *myo*-inositol. Structural elaborations on the membrane-oriented hydrophobic surface and a general expansion of PtdIns-binding pocket are highlighted in yellow (top row, center panel: surface rendering of the binding pocket; bottom row, center panel: ribbon representation of the aligned structures). HEK293-AT1 cells (10 μm scale bars) are shown expressing either the *Listeria* mRFP-PI-PLC (bottom row, left panel) or EGFP-*Bc*PI-PLC (bottom row, right panel). Please note the clear morphological changes associated with cells expressing the highly active EGFP-*Bc*PI-PLC, which are rounded and lack well-defined membranous organelles within the cytosol. (B) Detailed view of the *Bc*PI-PLC active site, highlighting the hydrogen bonding network involved in coordinating the inositol headgroup (dashed yellow lines) as well as the residues targeted for mutagenesis throughout this study. In particular, although His32 and His82 are both required for catalysis, notice that only His32 is directly involved in substrate coordination. A comparison of the localization of the (C) EGFP-*Bc*PI-PLC^H32A^ and (D) EGFP-*Bc*PI-PLC^H82A^ mutants are shown in HEK293-AT1 cells. Relative to the wild-type EGFP-*Bc*PI-PLC, both active site mutations rescue the drastic changes observed in the overall cell shape, but the EGFP-*Bc*PI-PLC^H82A^ construct clearly shows enhanced membrane binding relative to the EGFP-*Bc*PI-PLC^H32A^ mutant.

**Figure 2.**
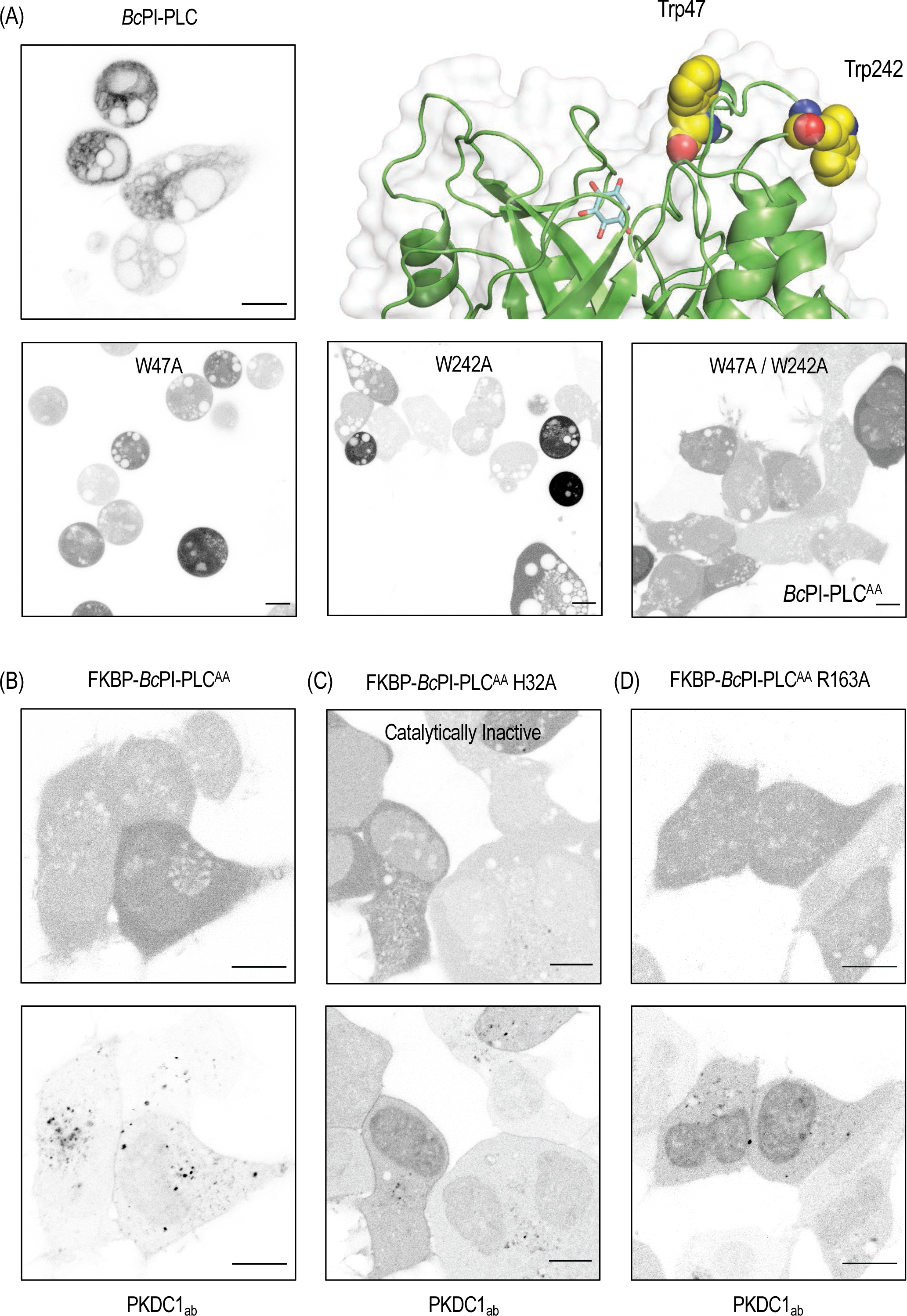
Mutagenesis of the *Bc*PI-PLC interface limits cytosolic activity by limiting the access to membrane-embedded substrate. (A) Mutagenesis of hydrophobic residues on the membrane-oriented surface were designed to disrupt the interfacial penetration of the *Bc*PI-PLC scaffold. A detailed view of the *Bc*PI-PLC interface is shown (top row, right) with residues Trp47 and Trp242 highlighted (atomic coloring, yellow spheres; PDB Accession: 1PTG; Heinz et al., 1995). Images from HEK293-AT1 cells (10 μm scale bars) show expression of the wild-type EGFP-*Bc*PI-PLC (low expression; top row, left panel) as well as the EGFP-*Bc*PI-PLC^W47A^ (bottom row, left panel), EGFP-*Bc*PI-PLC^W242A^ (bottom row, center panel), and EGFP-*Bc*PI-PLC^W47A/W242A^ (*Bc*PI-PLC^AA^; bottom row, right panel) mutants. Compared with the wild-type EGFP-*Bc*PI-PLC, the single tryptophan mutants are completely cytosolic, but still severely alter the cell morphology. However, expression of the EGFP-*Bc*PI-PLC^AA^ double-mutant shows clear cytosolic localization and almost no gross alteration to the cell shape; which is indicative of a significant drop in catalytic activity. Now cytosolic, FKBP-tagging of the *Bc*PI-PLC^AA^ scaffold was done to explore the utility of this modified enzyme for acute recruitment to FRB-labelled membranes. Co-expression of the high-affinity DAG-binding probe (mRFP-PKDC1_ab_; bottom row), which should reveal any changes in the localization of the PI-PLC hydrolytic product, is shown in HEK293-AT1 cells (10 μm scale bars) with the (B, left panels) mRFP-FKBP-*Bc*PI-PLC^AA^ backbone as well as with mutations that would inactivate (C, center panels; H32A) or modulate (D, right panels; R163A) the catalytic activity. Relative to the inactive control, the fully active mRFP-FKBP-*Bc*PI-PLC^AA^ does show a change in the cytosolic fraction of the PKDC1_ab_ probe, as well as the formation of some bright internal puncta. However, mutagenesis of the active site to FKBP-*Bc*PI-PLC^AA^ R163A prevents the gross redistribution of the DAG sensor.

**Figure 3.**
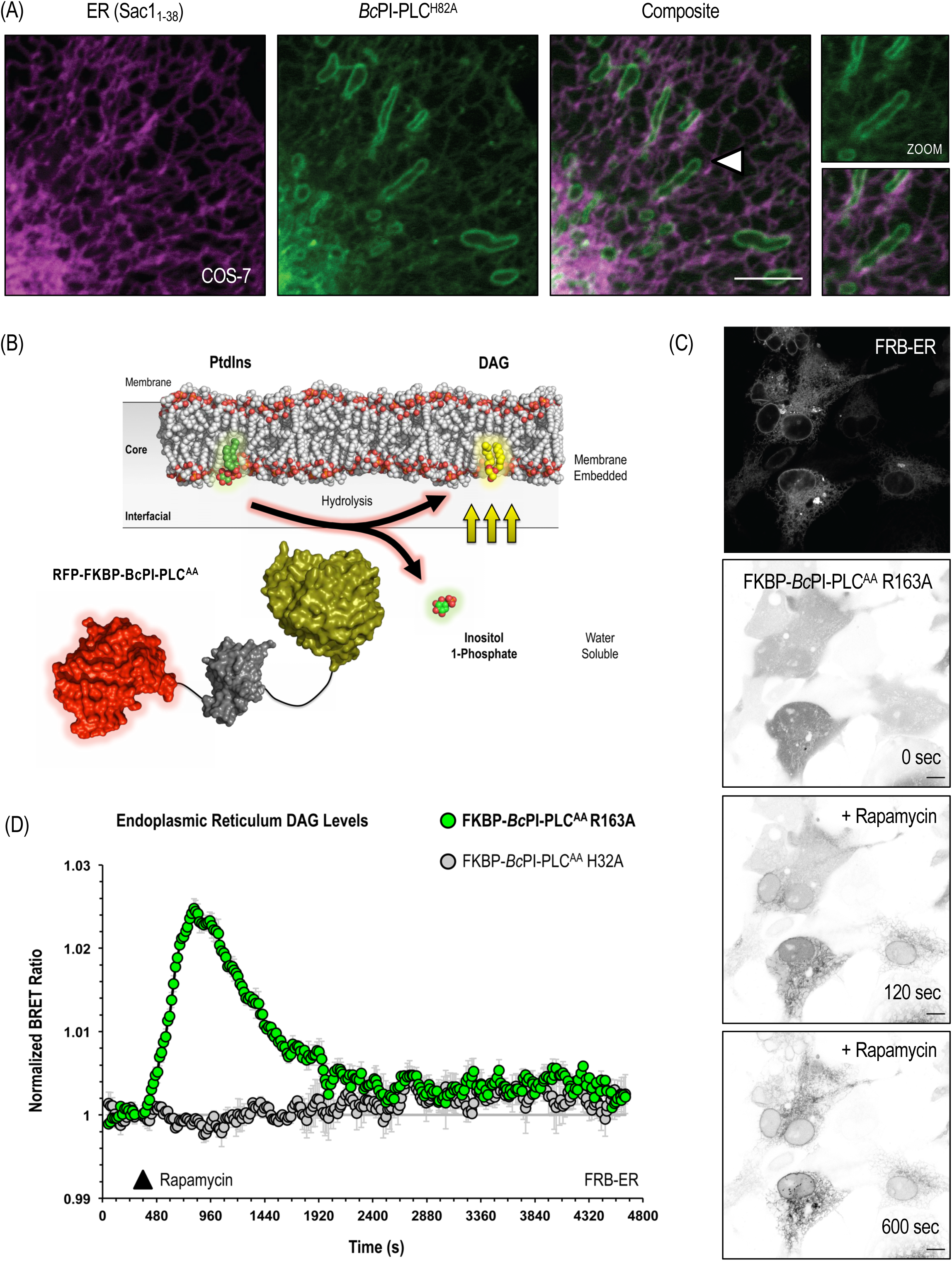
Acute manipulation of PtdIns content in the ER membrane. (A) As the biochemically-defined site of PtdIns synthesis, we examined the co-expression of EGFP-*Bc*PI-PLC^H82A^ and an ER marker, consisting of an mRFP-tagged signal sequence from the ER-resident protein Sac1 (mRFP-Sac1_1-38_), in COS-7 cells (5 μm scale bar). An enlarged view of the region identified by the white arrowhead not only shows the localization of EGFP-*Bc*PI-PLC^H82A^ to the ER tubules, but also highlights the relative enrichment of this probe within the outer mitochondrial membrane and at ER-mitochondria contacts. (B) Schematic depicting rapamycin-induced recruitment of FKBP-*Bc*PI-PLC^AA^ to locally hydrolyze PtdIns to produce membrane-embedded DAG and water soluble inositol 1-phosphate. In using this methodology, the local production of DAG can be used as a direct proxy for resting PtdIns content as well as provide information related to the turnover of PtdIns within a membrane compartment. (C) Time course detailing an example of rapamycin-induced recruitment of the mRFP-FKBP-*Bc*PI-PLC^AA^ R163A variant to the ER using the ER-FRB (mTagBFP2-FRB-CyB5_tail_) recruiter in HEK293-AT1 cells. (D) DAG production within the cytosolic leaflet of the ER was measured using the ER-DAG^BRET^ biosensor (sLuc-PKDC1_ab_-T2A-mVenus-CyB5A_tail_) in combination with the iRFP-FKBP-*Bc*PI-PLC^AA^ R163A and ER-FRB (mTagBFP2^E215A^-FRB-CyB5_tail_) heterodimerization system. Rapamycin (100 nM) was added manually after a 4 min baseline BRET measurement, with the post-rapamycin measurement beginning at ∼360s and continuing for 60 min (15 s / cycle). Recruitment of the iRFP-FKBP-*Bc*PI-PLC^AA^ R163A, but not the catalytically inactive iRFP-FKBP-*Bc*PI-PLC^AA^ H32A, resulted in the rapid accumulation of DAG within the ER; which was relatively short-lived and returned to basal levels after roughly 30 min. BRET measurements were carried out in triplicate wells (HEK293-AT1; 1×10^5^ cells/well) and repeated in three independent experiments. BRET ratios were first expressed relative to the baseline BRET measurement and then rapamycin-treated wells were normalized to the DMSO-treated controls.

**Figure 4.**
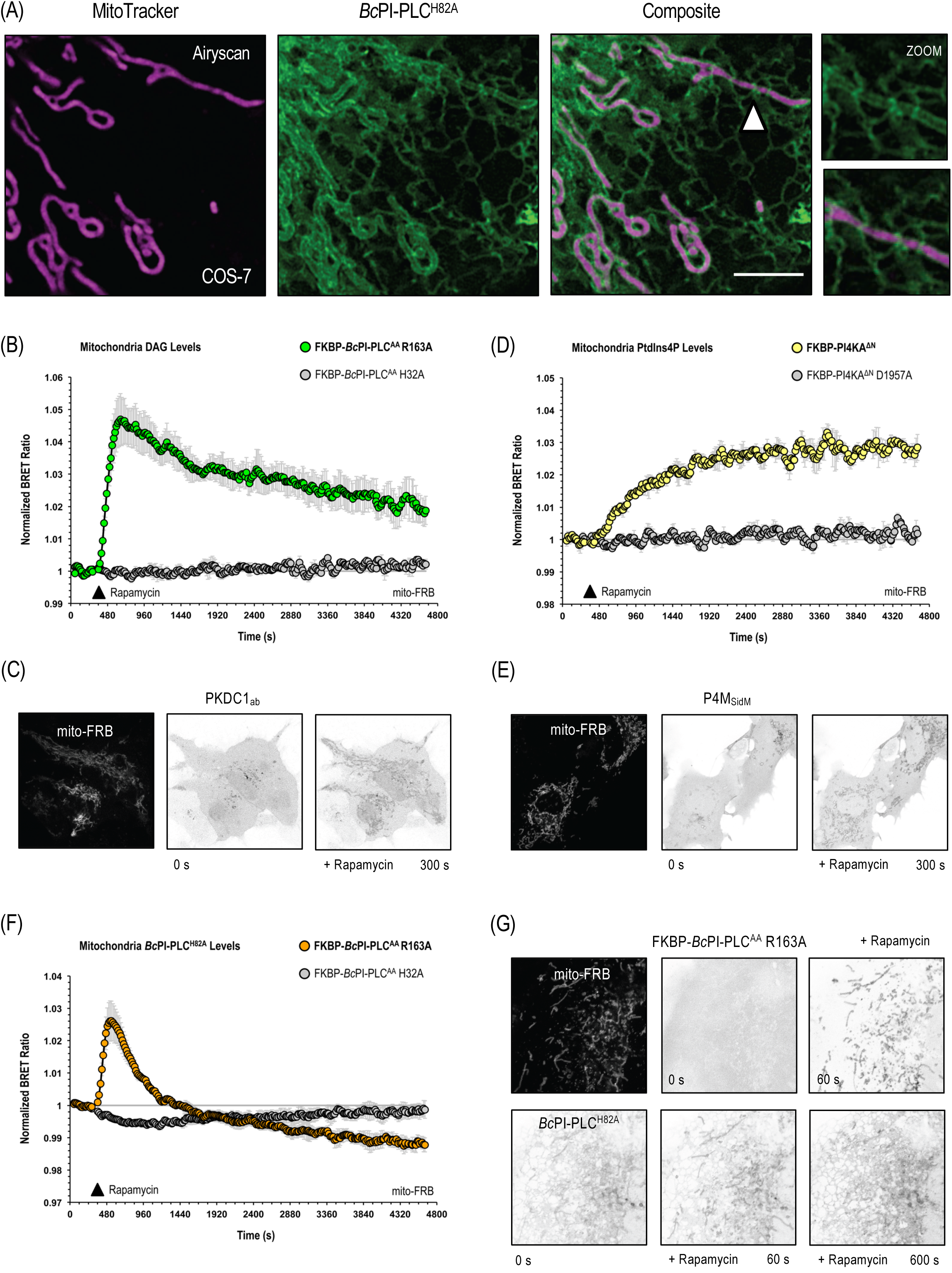
PtdIns is enriched in the outer mitochondrial membrane. (A) Localization of EGFP-*Bc*PI-PLC^H82A^ is shown in COS-7 cells loaded with the lumenal mitochondrial dye, MitoTracker, using the super-resolution Airyscan detector (5 μm scale bar). An enlarged view of the region identified by the white arrowhead shows the clear localization of EGFP-*Bc*PI-PLC^H82A^ to the membrane surrounding the mitochondrial lumen and further highlights the enrichment of the probe at ER-mitochondria contacts. (B) DAG production within the outer mitochondrial membrane was measured using the mito-DAG^BRET^ biosensor (AKAP-mVenus-T2A-sLuc-PKDC1_ab_) in combination with the iRFP-FKBP-*Bc*PI-PLC^AA^ R163A and mito-FRB (AKAP-FRB-ECFP^W66A^) heterodimerization system. (C) Representative images showing the mitochondrial recruiter (mito-FRB, AKAP-FRB-ECFP; left panel) as well as the initial (center panel) and post-rapamycin (100 nM; right panel) localization of a selective DAG-binding probe (EGFP-PKDC1_ab_). Together, these data show the rapid accumulation of DAG within the outer membrane of the mitochondria following the acute recruitment of FKBP-*Bc*PI-PLC^AA^ R163A. (D) PtdIns4P production within the outer mitochondrial membrane was measured using the mito-PtdIns4P^BRET^ biosensor (AKAP-mVenus-T2A-sLuc-P4M_SidM_^X2^) in combination with the iRFP-FKBP-PI4KA^ΔN^ and mito-FRB (AKAP-FRB-ECFP^W66A^) heterodimerization system. (E) Representative images showing the mitochondrial recruiter (AKAP-FRB-ECFP; left panel) as well as the initial (center panel) and post-rapamycin (100 nM; right panel) localization of a selective PtdIns4P-binding probe (EGFP-P4M_SidM_). The rapid accumulation of PtdIns4P following the acute recruitment of iRFP-FKBP-PI4KA^ΔN^ to the surface of the mitochondria further supports the conclusion that PtdIns is metabolically available within the outer mitochondrial membrane. (F) *Bc*PI-PLC^H82A^ localization to the outer mitochondrial membrane was measured using the mito-H82A^BRET^ biosensor (AKAP-mVenus-T2A-sLuc-*Bc*PI-PLC^H82A^) in combination with the iRFP-FKBP-*Bc*PI-PLC^AA^ R163A and mito-FRB (AKAP-FRB-ECFP^W66A^) heterodimerization system. After an initial accumulation of the probe, which could be related to compensatory changes in lipid transfer or reflect the introduction of local packing defects, *Bc*PI-PLC^H82A^ is rapidly lost from the surface of the mitochondria and levels eventually cross below the pre-treatment values. (G) Representative images showing the mitochondrial recruiter (mito-FRB, AKAP-FRB-ECFP; top row, left panel) and the rapid rapamycin-induced (100 nM) recruitment of the mRFP-FKBP-*Bc*PI-PLC^AA^ R163A enzyme to the outer mitochondrial membrane (top row, center and right panels). The still images also show the transient accumulation (60s) and eventual loss (600s) of the EGFP-*Bc*PI-PLC^H82A^ probe from the surface of the mitochondria. Please note that for each of the membrane recruitment studies using BRET-based measurements (B, D, and F), rapamycin (100 nM) was added manually after a 4 min baseline BRET measurement; with the post-rapamycin measurement beginning at ∼360s and continuing for 60 min (15 s / cycle). All measurements were carried out in triplicate wells (HEK293-AT1; 1×10^5^ cells/well) and repeated in three independent experiments. BRET ratios were first expressed relative to the baseline BRET measurement and then rapamycin-treated wells were normalized to the DMSO-treated controls.

**Figure 5.**
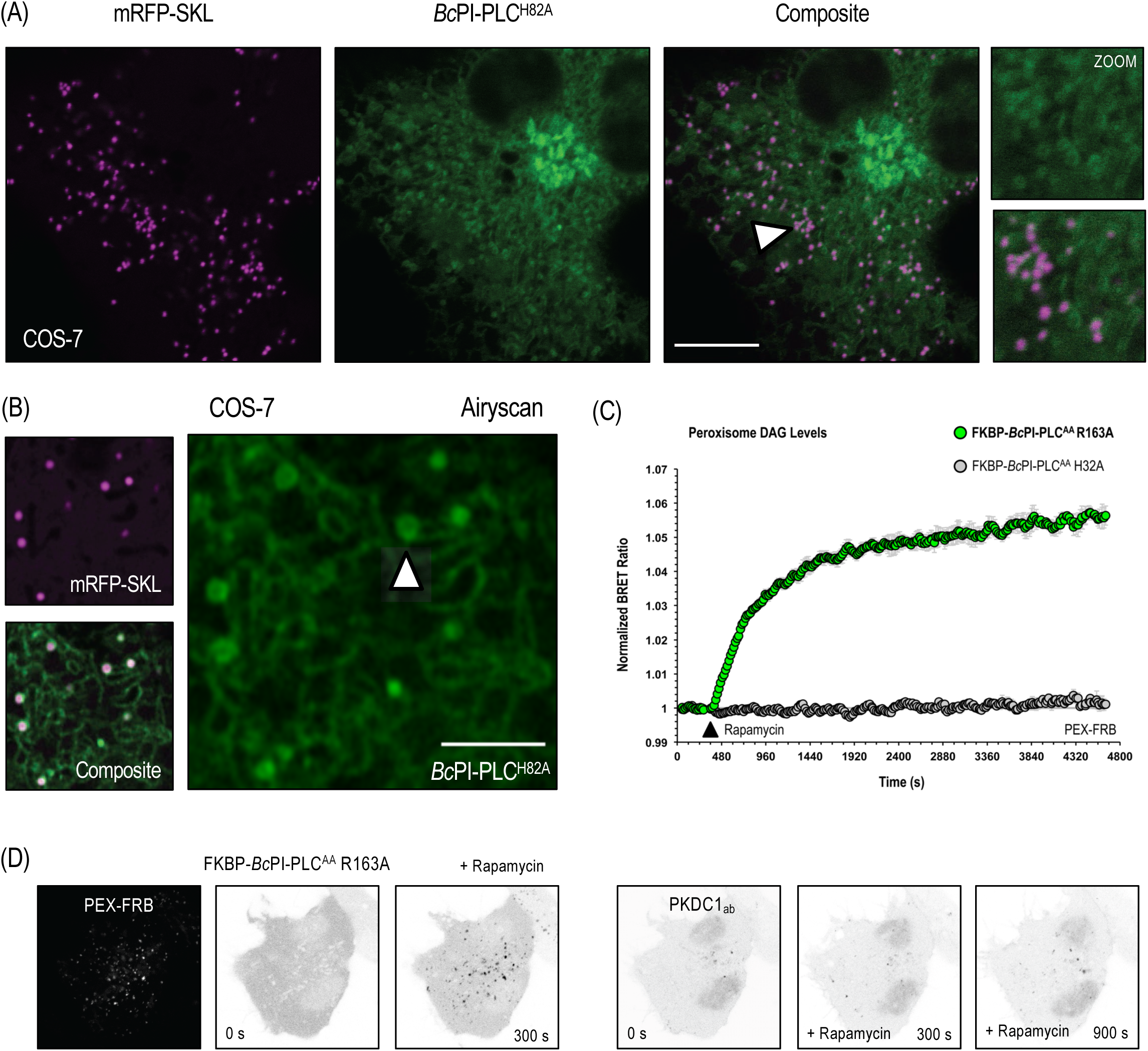
PtdIns is enriched in the cytosolic leaflet of peroxisomes. (A) Co-expression of EGFP-*Bc*PI-PLC^H82A^ with the lumenally-targeted mRFP-SKL peroxisome marker, which seems to enlarge the peroxisome compartment, in COS-7 cells (10 μm scale bar). An enlarged view of the region identified by the white arrowhead shows the wrapping of the EGFP-*Bc*PI-PLC^H82A^ around the punctate mRFP-SKL core. (B) Localization of EGFP-*Bc*PI-PLC^H82A^ in COS-7 cells co-expressing mRFP-SKL using the super-resolution Airyscan detector (2.5 μm scale bar). An enlarged view of the EGFP-*Bc*PI-PLC^H82A^ channel is provided to highlight the ability to resolve the hollowed core of the signal from the EGFP-*Bc*PI-PLC^H82A^, which appears to envelope the peroxisome puncta (white arrowhead). (C) DAG production within the cytosolic leaflet of the peroxisomes was measured using the PEX-DAG^BRET^ biosensor (PEX-mVenus-T2A-sLuc-PKDC1_ab_) in combination with the iRFP-FKBP-*Bc*PI-PLC^AA^ R163A and PEX-FRB (PEX3_1-42_-FRB-ECFP^W66A^) heterodimerization system. Rapamycin (100 nM) was added manually after a 4 min baseline BRET measurement, with the post-rapamycin measurement beginning at ∼360s and continuing for 60 min (15 s / cycle). Measurements were carried out in triplicate wells (HEK293-AT1; 1×10^5^ cells/well) and repeated in three independent experiments. BRET ratios were first expressed relative to the baseline BRET measurement and then rapamycin-treated wells were normalized to the DMSO-treated controls. (D) Representative images showing the distribution of the peroxisome recruiter (PEX-FRB, PEX3_1-42_-FRB-ECFP) and the rapid rapamycin-induced (100 nM) recruitment of the mRFP-FKBP-*Bc*PI-PLC^AA^ R163A enzyme to the peroxisomes (left panels). Overall, the BRET-based measurements show a rapid and sustained accumulation of DAG on the surface of peroxisomes following the acute recruitment of FKBP-*Bc*PI-PLC^AA^ R163A; however, only minor changes in the localization of the DAG probe (EGFP-PKDC1_ab_; right panels), including the transient formation of cytosolic puncta, were seen by confocal microscopy using HEK293-AT1 cells.

**Figure 6.**
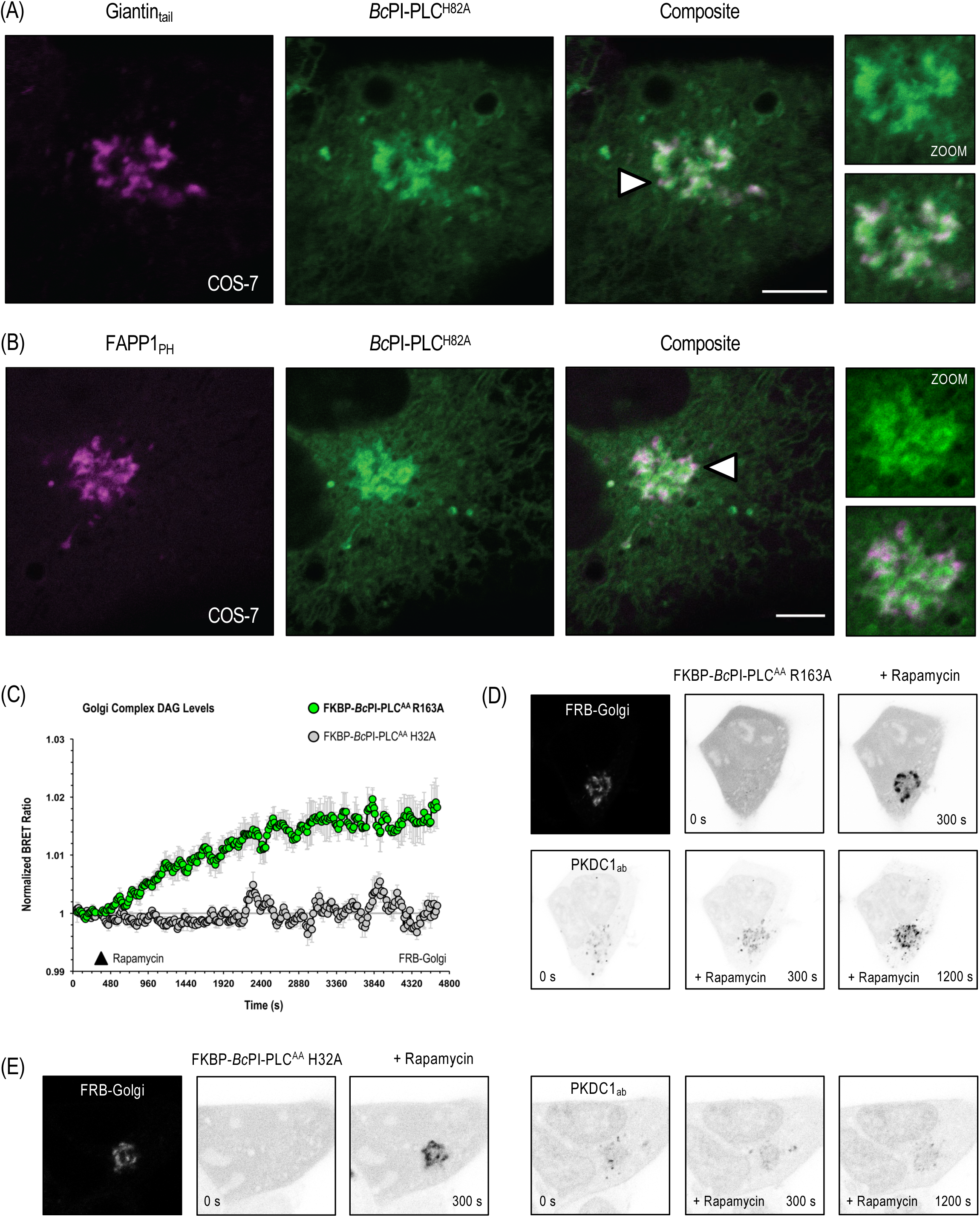
PtdIns is enriched at the Golgi Complex. Co-expression of EGFP-*Bc*PI-PLC^H82A^ in COS-7 cells (5 μm scale bar) with an integral Golgi-localized membrane protein (A; FRB-mCherry-Giantin_tail_) or the Arf1- and PtdIns4P-sensitive pleckstrin homology (PH) domain of FAPP1 (B; FAPP1_PH_-EGFP). An enlarged view of the regions identified by the white arrowheads not only shows the clear co-localization of EGFP-*Bc*PI-PLC^H82A^ with markers of the Golgi complex, but also highlights the specific enrichment of these labels, sometimes independent of one another, within the perinuclear Golgi region. Of particular interest is the concentration of the FAPP1_PH_ domain within peripheral puncta at the edges of the Golgi complex in regions that appear to be distinct from those labeled with the EGFP-*Bc*PI-PLC^H82A^ probe. (C) DAG production within the Golgi complex was measured using the Golgi-DAG^BRET^ biosensor (sLuc-PKDC1_ab_-T2A-mVenus-Giantin_tail_) in combination with the iRFP-FKBP-*Bc*PI-PLC^AA^ R163A and FRB-Golgi (ECFP^W66A^-FRB-Giantin_tail_) heterodimerization system. Rapamycin (100 nM) was added manually after a 4 min baseline BRET measurement, with the post-rapamycin measurement beginning at ∼360s and continuing for 60 min (15 s / cycle). Recruitment of the catalytically active iRFP-FKBP-*Bc*PI-PLC^AA^ R163A, but not the inactive iRFP-FKBP-*Bc*PI-PLC^AA^ H32A, resulted in the sustained accumulation of DAG within the cytosolic leaflet of the Golgi complex. All measurements were carried out in triplicate wells (HEK293-AT1; 1×10^5^ cells/well) and repeated in three independent experiments. BRET ratios were first expressed relative to the baseline BRET measurement and then rapamycin-treated wells were normalized to the DMSO-treated controls. (D) Representative images showing the Golgi membrane recruiter (FRB-Golgi, ECFP-FRB-Giantin_tail_) and the rapamycin-induced (100 nM) recruitment of the mRFP-FKBP-*Bc*PI-PLC^AA^ R163A enzyme to the Golgi complex (top row). The still images show the sustained accumulation of DAG (EGFP-PKDC1_ab_) on the surface of the Golgi complex (bottom row). (E) Importantly, as shown in the representative images, recruitment of the inactive enzyme (FKBP-*Bc*PI-PLC^AA^ H32A) to the Golgi complex (left panels) did not alter the localization of the EGFP-PKDC1_ab_ probe (right panels).

### Acute Manipulation of PtdIns in defined membrane compartments

To investigate the effects of spatially restricted PtdIns hydrolysis in specific membrane compartments, we took advantage of the chemically-inducible protein heterodimerization system that uses rapamycin-dependent association of FK506-binding protein (FKBP) with the FKBP-rapamycin binding (FRB) domain from mammalian target of rapamycin (Choi et al., 1996; Liang et al., 1999). This system was exploited to recruit a modified, but catalytically-active, FKBP-*Bc*PI-PLC to specific organelle membranes marked with FRB-conjugated targeting proteins (Belshaw et al., 1996; Komatsu et al., 2010). This approach has been successfully applied for other lipid-modifying enzymes (Varnai et al., 2006; Fili et al., 2006; Suh et al., 2006; Heo et al., 2006; Szentpetery et al., 2010; Hammond et al., 2012) and using them in concert with the *Bc*PI-PLC would provide a powerful new strategy to map the membrane distributions of PtdIns. Importantly, our previous effort to generate a recruitable PI-PLC construct using the *Listeria monocytogenes* enzyme (Kim et al., 2011) did not yield a generally useful tool in our hands. Briefly, poor membrane recruitment combined with the relatively limited catalytic activity of the *Listeria* PI-PLC enzyme were quickly identified as the main reasons for its limited utility. Therefore, we again turned to the *Bc*PI-PLC and wanted to generate mutants with reduced membrane-binding to decrease their background hydrolysis of PtdIns from the cytosol. Recruiting these enzyme variants to target membranes, in close proximity to PtdIns, with the rapamycin-inducible heterodimerization system would be useful as an alternative strategy to map PtdIns distributions in intact cells.

With these goals in mind, we examined the *Bc*PI-PLC crystal structure, as well as the existing *in vitro* binding studies, to find any exposed hydrophobic or charged residues on the membrane-oriented surface of the enzyme that were likely to contribute to interfacial binding. We identified two membrane-oriented tryptophan residues, namely Trp47 within helix B and Trp242 in the extended α7-βG loop (Figure 2a), facing which have already been shown to facilitate the association of *Bacillus* PI-PLCs with phospholipid vesicles (Feng et al., 2002; Guo et al., 2008; Khan et al., 2016). Aromatic or hydrophobic side chains, and tryptophan residues in particular, are known to reinforce the membrane attachment of peripheral proteins by directly penetrating into the hydrocarbon core of the bilayer (Yau et al., 1998; Lomize et al., 2007; Fuglebakk and Reuter, 2018). Given the importance of membrane-binding events for access to lipid substrates, the catalytic activity of many phospholipase families display marked interfacial activation and the *Bacillus* PI-PLCs, specifically, exhibit a substantial preference for substrates presented within an interface rather than as monomers in solution (Lewis et al., 1993; Zhou et al., 1997a,b; Qian et al., 1998). Expression of *Bc*PI-PLC with point mutations of either Trp47 (W47A) or Trp242 (W242A) already failed to strongly interact with subcellular membranes and were almost completely cytosolic; but, even without strong membrane association, the cells expressing either of these constructs still suffered from marked cytotoxicity (Fig.2A). However, mutagenesis of both Trp47 and Trp242 (W47A/W242A) prevented the membrane binding of the *Bc*PI-PLC and also mitigated the cytotoxicity (Fig.2A). Now cytosolic, we could tag the *Bc*PI-PLC^W47A/W242A^ mutant with the FKBP dimerization module to create a chimeric enzyme construct (FKBP-*Bc*PI-PLC^AA^) that could be rapidly recruited to distinct organelle membranes with high spatial and temporal resolution (Fig.S3A). Importantly, relative to the inactive FKBP-*Bc*PI-PLC^AA^ H32A variant, we did not observe drastic alterations to the distribution of an established biosensor for the PtdIns hydrolytic product, DAG (GFP-PKDC1_ab_; Kim et al., 2011), in cells with low to moderate expression of FKBP-*Bc*PI-PLC^AA^ (Fig.2B and 2C); supporting the conclusion that interfacial mutations of the *Bc*PI-PLC effectively reduced the catalytic activity of this enzymes from the cytosol. However, we noted that cells expressing high levels of the FKBP-*Bc*PI-PLC^AA^ also showed a reduced nuclear fraction as well as punctate endomembrane membrane localization of the high-affinity DAG probe, suggesting that this modified enzyme still retained some basal catalytic activity. Therefore, we decided to further modify the FKBP-*Bc*PI-PLC^AA^ backbone by using mutagenesis of the enzyme active site in an attempt to further limit enzymatic activity from the cytosol to optimize the dynamic range of the induced catalysis. After excluding mutations associated with excessive reductions in catalytic activity *in vitro* (Gässler et al., 1997), we chose residues with accessory roles related to substrate binding or coordination; specifically targeting Arg163 and Tyr200 (Fig.1B). However, in order to compare the relative activities of the recruitable *Bc*PI-PLC variants and account for individual cell variability, we wanted to address the problem of detecting localized DAG content, the lipid product associated with their activity, at the level of entire cell populations. For this, we worked to establish a semi-high throughput approach for the quantification of changes in subcellular membrane lipid compositions that complements the established single cell analyses that rely on confocal microscopy.

### BRET-based biosensors allow for population-level analyses of lipid dynamics within defined membrane compartments using live cells

To allow for comparative measurements of localized *Bc*PI-PLC activity in whole populations of live cells, we created a series of BRET-based biosensors that measure localized DAG levels by monitoring the resonance enery transfer between a high-affinity DAG probe and organelle-specific membrane labels. This methodology relies on a single-plasmid design that incorporates the self-cleaving viral 2A peptide from *Thosea asigna* (T2A; Donnelly et al., 2001; Szymczak et al., 2004; Liu et al., 2017). The internal T2A site facilitates the split of the translated polypeptide into two proteins and allows for the uniform and roughly equimolar expression of both the organelle-anchored BRET acceptor (mVenus) and Luciferase-tagged lipid-binding probe, which serves as the BRET donor (Varnai et al., 2017). Due to the prominent localization of the *Bc*PI-PLC^H82A^ probe within the outer mitochondrial membrane, we chose to validate the design of the BRET-based biosensors by attempting to monitor mitochondrial DAG levels. Briefly, the mitochondrial targeting sequence of AKAP (Csordás et al., 2010) was fused to mVenus, while sLuc was fused to the PKDC1_ab_ probe to produce the compartment-selective mito-DAG^BRET^ biosensor (AKAP-mVenus-T2A-sLuc-PKDC1_ab_). The mito-DAG^BRET^reporter was then used in combination with a mitochondria-targeted FRB construct (mito-FRB; AKAP-FRB-ECFP) and the cytosolic FKBP-*Bc*PI-PLC^AA^ enzyme variant to induce PtdIns hydrolysis on the surface of mitochondria. Importantly, to prevent artefacts from the enzyme recruitment, which could interfere with the energy transfer process, the chromophore of the ECFP (W66A; Heim et al., 1994) or mTagBFP2 (E215A; Subach et al., 2011) fluorescent proteins that are present within the membrane-targeting FRB constructs were disrupted by mutagenesis; while the FKBP-*Bc*PI-PLC^AA^ construct was tagged with the BRET-compatible lablel, iRFP. By changing either the membrane targeting motif or the lipid-binding reporter present within the vector backbone, we were able to create a variety of biosensors to monitor changes in various lipid species within specific membrane compartments of live cells.

Initial results using the FKBP-*Bc*PI-PLC^AA^ recruitment system with the mito-DAG^BRET^ biosensor demonstrated the rapid production of DAG within the outer mitochondrial membrane following acute treatment with rapamycin (100 nM; Fig.S3B and S3C). Importantly, for these and all remaining studies, recruitment of an inactive enzyme variant, FKBP-*Bc*PI-PLC^AA^ H32A, was used as a transfection control and for the normalization of the BRET data. Overall, in establishing this experimental strategy, the acute production of DAG by the recruited FKBP-*Bc*PI-PLC^AA^ served as an indirect proxy of PtdIns availability within intact cell membranes and studies using this approach were then applied to other organelle membranes in order to validate the PtdIns distributions mapped using the *Bc*PI-PLC^H82A^ probe. In an attempt to enhance the dynamic range of the recruited enzyme construct, we generated additional mutations in the active site of the FKBP-*Bc*PI-PLC^AA^ scaffold to reduce its enzymatic activity. Relative to the parent FKBP-*Bc*PI-PLC^AA^ construct, alanine substitution or conservative mutagenesis of Arg163 (R163A and R163K; Fig.S3B) or Tyr200 (Y200A and Y200F; Fig.S3C) showed clear alterations in the kinetics of the DAG production observed after enzyme recruitment. The minor elevation of the basal mito-DAG^BRET^ signals, which are not seen in the normalized curves (Fig. S3C), as well as the rapid initial peak in the DAG signal observed for the FKBP-*Bc*PI-PLC^AA^ R163A, R163K, and Y200F variants suggested a minimized consumption and better preservation of the basal PtdIns levels by the mutant enzymes before membrane recruitment; features that established these enzyme variants as ideal candidates for further characterization. Subsequently, after examining the basal distributions of the DAG sensor (Fig.2D), as well as repeating the mitochondrial recruitment experiments using single-cell confocal microscopy, we chose the FKBP-*Bc*PI-PLC^AA^ R163A variant for the remaining studies due to the clear benefits of the enhanced signal to noise ratio for enzyme recruitment assays. Overall, these efforts using iterative mutagenesis have produced a series of FKBP-*Bc*PI-PLC^AA^ constructs with a range of catalytic activities that can be selected for investigating diverse questions related to the availability and turnover of membrane PtdIns.

### PtdIns is detected in the cytosolic membrane leaflets of the ER, mitochondria, and peroxisomes

The presence of the synthetic machinery responsible for PtdIns production, as well as the results from early membrane fractionation studies (Antonsson, 1997; Vance, 2015), suggested that PtdIns is likely to be an important component of ER membranes. Expression of the GFP-*Bc*PI-PLC^H82A^ probe with an ER-marker (mRFP-ER(Sac1_1-38_); Várnai et al., 2007) show clear co-localization within the ER tubules and the perinuclear region (COS-7 cells; Fig.3A). As a further proof of the presence of PtdIns within the ER membrane, we used FKBP-*Bc*PI-PLC^AA^ R163A and an ER-targeted FRB (FRB-ER; mTagBFP2-FRB-CyB5A_tail_) to acutely recruit the active enzyme to the surface of the ER. Local production of DAG within the cytosolic leaflet of the ER was measured using a novel ER-DAG^BRET^ biosensor (sLuc-PKDC1_ab_-T2A-mVenus-CyB5A_tail_; Fig.3B and 3C). Recruitment of the FKBP-*Bc*PI-PLC^AA^ R163A resulted in the rapid accumulation of DAG within the ER, which was relatively short-lived and quickly returned to basal levels after roughly 30 min (Fig.3D). The transient nature of the DAG production observed is likely to reflect the active utilization of the DAG produced for other metabolic functions within the ER. In addition to labeling the tubular ER, the outer membrane of the mitochondria showed a clear enrichment of the *Bc*PI-PLC^H82A^ probe. In particular, although the mitochondrial localization of *Bc*PI-PLC^H82A^ could be seen when co-expressing fluorescent markers of the ER (Fig.4A), but was further enhanced using super-resolution imaging (Airyscan; Huff, 2015; Scipioni et al., 2018) and the lumenal mitochondrial dye, MitoTracker (Chazotte, 2011; Fig.4B). These high-resolution images also highlight an apparent concentration of the *Bc*PI-PLC^H82A^ probe at the ER-mitochondria interface; including at contacts where the ER appears to wrap around the mitochondrial membrane (Fig.4A; white arrowhead). During these imaging studies, we also observed *Bc*PI-PLC^H82A^ localization to small rounded structures, often associated with the mitochondria or ER, that did not appear to be part of the Golgi complex or endosomal system. Co-expression of GFP-*Bc*PI-PLC^H82A^ with a fluorescently-tagged protein containing a peroxisomal targeting signal (mRFP-SKL; Kim et al., 2006) confirmed the identity of these membrane compartments as peroxisomes; with the *Bc*PI-PLC^H82A^ actually enveloping the lumenally-targeted fluorophore (Fig.5A and 5B; white arrowheads). In agreement with these localization data, rapamycin-dependent recruitment of FKBP-*Bc*PI-PLC^AA^ R163A caused a rapid and sustained increase of DAG within the outer mitochondrial membrane (mito-DAG^BRET^; Fig.4B and 4C) and peroxisomes (PEX-DAG^BRET^, PEX-mVenus-T2A-sLuc-PKDC1_ab_; Fig.5C and 5D). Given the mitochondrial enrichment based on the localization of the *Bc*PI-PLC^H82A^ probe, we looked for additional methods to evaluate the presence of PtdIns in the mitochondrial outer membrane. Specifically, recruitment of a truncated variant of a PtdIns 4-kinase (FKBP-PI4KA^ΔN^; Hammond et al., 2014) elicited the local production of PtdIns4P on the mitochondrial surface, as measured using a mito-PI4P^BRET^ biosensor (AKAP-mVenus-T2A-sLuc-P4M_SidM_^x2^; Fig.4D and 4E). Taken together, these data support the conclusion that PtdIns is localized within the outer mitochondrial membrane and accessible to cytosolic effectors.

Given the strong evidence for the specific enrichment of PtdIns within the outer mitochondrial membrane, we next used the recruitable FKBP-*Bc*PI-PLC^AA^ R163A enzyme to try and verify whether the membrane localization of the *Bc*PI-PLC^H82A^ probe responded to the hydrolysis of PtdIns on the surface of mitochondria. Surprisingly, recruitment of FKBP-*Bc*PI-PLC^AA^ R163A to the mitochondria caused a rapid increase in the association of *Bc*PI-PLC^H82A^ (mito-H82A^BRET^, AKAP-mVenus-T2A-sLuc-*Bc*PI-PLC^H82A^) with the outer mitochondrial membrane. This initial increase, however, was followed by the steady elimination of the *Bc*PI-PLC^H82A^ probe from the surface of the mitochondria (Fig.4F and 4G). This redistribution could reflect an increased insertion of the probe into the membrane in response to the localized introduction of membrane packing defects. In fact, a recent biophysical study has shown that the catalytic activity of the *Bc*PI-PLC is enhanced by the accumulation of the enzymatic product, DAG, which appears to function by locally decreasing membrane order to facilitate interfacial binding (Ahyayauch et al., 2015). Importantly, however, simple production of DAG is not sufficient to drive the membrane localization of the *Bc*PI-PLC^H82A^ probe (see below; Fig.9I).Taken together, these data highlight the metabolic availability of PtdIns within the outer mitochondrial membrane and, in carrying out these experiments, it did not escape our attention that acute hydrolysis of the mitochondrial PtdIns pool, as well as the associated local production of DAG, caused profound changes to the structure of the mitochondrial network. The ability to alter mitochondrial dynamics by initiating acute changes in membrane lipid composition represents an exciting application of these molecular tools that is being actively pursued in our laboratory.

### PtdIns is enriched in membranes of the Golgi complex and supports local PtdIns4P production

Along with the inner leaflet of the PM, the membranes of the Golgi complex show a significant enrichment with PtdIns4P, which is produced primarily by the type III PI 4-kinase (PI4K), PI4KB (Godi et al., 1999), with additional contributions from both type II PI4K enzymes, PI4K2A and PI4K2B (Wang et al., 2003; Weixel et al., 2005; Boura and Nencka, 2015). The production of PtdIns4P at the Golgi complex must be tightly regulated since it is an essential component of membrane trafficking systems (Wang et al., 2003; Godi et al., 2004) and also integrates with PPIn-dependent turnover throughout the endosomal system (Balla et al., 2002; Minogue et al., 2006; Dong et al., 2016; Baba et al., 2019). Co-localization of *Bc*PI-PLC^H82A^ with either the integral Golgi-localized membrane protein, Giantin, or the Arf1- and PtdIns4P-sensitive pleckstrin homology (PH) domain of FAPP1 (FAPP1_PH_; Levine and Munro, 2002; Szentpetery et al., 2010), showed clear overlap at the Golgi complex (Fig.6A and 6B). However, both markers also localized to peripheral areas of the perinuclear Golgi region that did not contain the *Bc*PI-PLC^H82A^ (Fig.6A and 6B, white arrowheads). Recruitment studies using FKBP-*Bc*PI-PLC^AA^ R163A and a Golgi-localized FRB-Giantin revealed a gradual and sustained increase of DAG (Golgi-DAG^BRET^, sLuc-PKDC1_ab_-T2A-mVenus-Giantin) after acute treatment with rapamycin that did not occur with the recruitment of the inactive control (FKBP-*Bc*PI-PLC^AA^ H32A; Fig.6C, 6D, and 6E). The kinetics of the DAG increase did not have the same rapid onset as those observed in the ER, outer mitochondrial membrane, or peroxisomes. This could reflect the fact that, relative to the amount of

DAG produced by the recruited enzyme, the steady-state levels of DAG are already high in the Golgi complex (Litvak et al., 2005). Next, to examine the localized dynamics of PtdIns turnover and to verify that the *Bc*PI-PLC^H82A^ probe indeed reflects an enrichment of PtdIns at the Golgi complex, we monitored the localization of GFP-*Bc*PI-PLC^H82A^ at the Golgi after the recruitment of FKBP-*Bc*PI-PLC^AA^ variants either directly to the surface of the Golgi complex or to the ER. Direct recruitment of FKBP-*Bc*PI-PLC^AA^ or the R163A mutant to the Golgi had a highly variable effect on the localization of the GFP-*Bc*PI-PLC^H82A^ probe at the Golgi complex: many cells showed a gradual, but almost complete loss of Golgi-localized GFP-*Bc*PI-PLC^H82A^ when recruiting the more active FKBP-*Bc*PI-PLC^AA^ enzyme directly to the surface of the Golgi (Fig.7A and 7B). However, other cells showed an increased localization of GFP-*Bc*PI-PLC^H82A^ probe similar to what was observed in the mitochondria. Again, this effect could be related to the increase in local packing defects generated in the Golgi membranes, by acute hydrolysis of resident PtdIns. Overall, pooled averages from live-cell imaging experiments monitoring the Golgi-specific recruitment of the FKBP-*Bc*PI-PLC^AA^ variants showed a relative increase and no clear depletion of PtdIns based on the localization of the GFP-*Bc*PI-PLC^H82A^ probe (Fig.7A). The relative inability to rapidly deplete PtdIns levels at the Golgi complex likely reflects the larger size of the steady-state PtdIns pool within this compartment and also suggest an efficient supply of newly-synthesized PtdIns from the ER. In line with this latter hypothesis, the recruitment of either FKBP-*Bc*PI-PLC^AA^ or the R163A variant to the ER, rather than directly to the Golgi complex, resulted in a rapid loss of GFP-*Bc*PI-PLC^H82A^ from the Golgi region (Fig.7C and 7D). Taken together, these data reveal a high capacity for PtdIns transport from the ER to the Golgi complex for the maintenance of the steady state levels of PtdIns.

**Figure 7.**
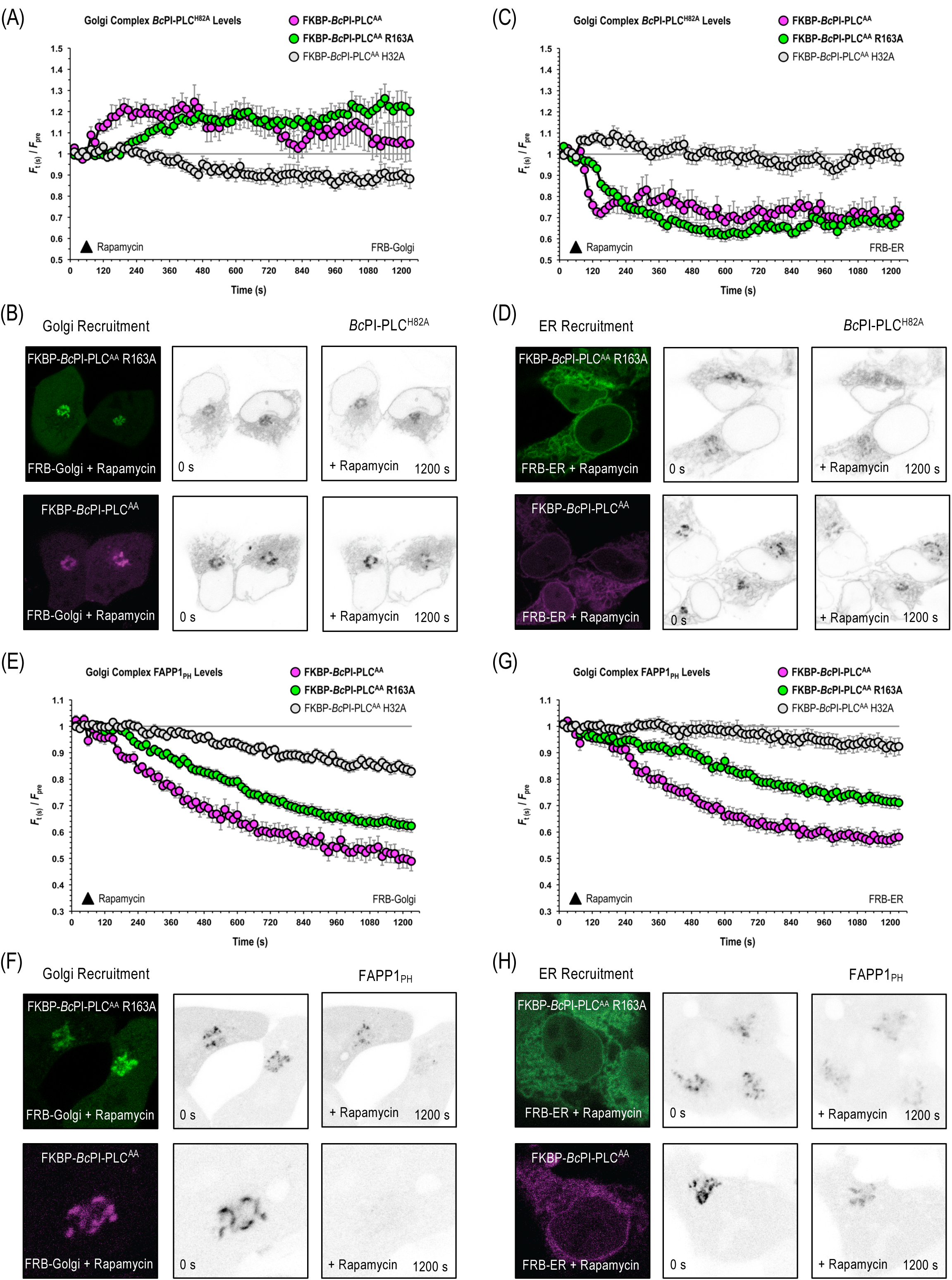
PtdIns4P levels at the Golgi complex are sensitive to the local availability of PtdIns. Pooled image analyses of (A and C) EGFP-*Bc*PI-PLC^H82A^ or (E and G) FAPP1_PH_-EGFP levels in the perinuclear Golgi region following recruitment of mRFP-FKBP-*Bc*PI-PLC^AA^ R163A (37 cells, FRB-Golgi + EGFP-*Bc*PI-PLC^H82A^; 27 cells, FRB-ER + EGFP-*Bc*PI-PLC^H82A^; 46 cells, FRB-Golgi + FAPP1_PH_-EGFP; 42 cells, FRB-ER + FAPP1_PH_-EGFP), mRFP-FKBP-*Bc*PI-PLC^AA^ (14 cells, FRB-Golgi + EGFP-*Bc*PI-PLC^H82A^; 16 cells, FRB-ER + EGFP-*Bc*PI-PLC^H82A^; 16 cells, FRB-Golgi + FAPP1_PH_-EGFP; 21 cells, FRB-ER + FAPP1_PH_-EGFP), or mRFP-FKBP-*Bc*PI-PLC^AA^ H32A (32 cells, FRB-Golgi + EGFP-*Bc*PI-PLC^H82A^; 27 cells, FRB-ER + EGFP-*Bc*PI-PLC^H82A^; 43 cells, FRB-Golgi + FAPP1_PH_-EGFP; 42 cells, FRB-ER + FAPP1_PH_-EGFP) either directly to the surface of the Golgi complex (A and E; FRB-Golgi, ECFP-FRB-Giantin_tail_) or indirectly to the site of PtdIns synthesis in the ER (C and G; FRB-ER, mTagBFP2-FRB-CyB5_tail_). Overall, results show the rapid depletion of FAPP1_PH_-EGFP at the Golgi complex following the recruitment of either FKBP-*Bc*PI-PLC^AA^ or the R163A variant to both the Golgi complex or to the ER. Direct recruitment of the FKBP-*Bc*PI-PLC^AA^ variants to the Golgi did not deplete EGFP-*Bc*PI-PLC^H82A^ from the surface of the Golgi complex, however both the FKBP-*Bc*PI-PLC^AA^ and R163A variant were able to rapidly drop EGFP-*Bc*PI-PLC^H82A^ levels in the perinuclear Golgi region when recruited to the ER. For each treatment using the active FKBP-*Bc*PI-PLC^AA^ or R163A enzymes, images of representative cells are shown with the resulting post-rapamycin (100nM) enzyme recruitment (right panel) as well as the initial (center panel) and final (left panel) localization of the EGFP-*Bc*PI-PLC^H82A^ or FAPP1_PH_-EGFP probes following 20 min of spatially-restricted PtdIns hydrolysis. Alternatively, each quantified graph shows the normalized intensities (*F*(t)/*F*_pre_) of the EGFP-*Bc*PI-PLC^H82A^ or FAPP1_PH_-EGFP signals at the perinuclear Golgi region, shown relative to the cytosol, following treatment with rapamycin (100nM) using HEK293-AT1 cells. Data from these image analyses (A, C, E, and G) are presented as mean values ± SEM from a minimum of four independent experiment. The pre-treatment period used for normalization was defined as the average ratio of the Golgi:cytosolic signal measured over the first 4 frames of each recording, prior to rapamycin addition.

To examine how localized changes in PtdIns availability would affect the Golgi PtdIns4P levels, we recruited the FKBP-*Bc*PI-PLC^AA^ variants to the Golgi complex and monitored the localization of FAPP1_PH_-GFP. We observed that both direct recruitment of the FKBP-*Bc*PI-PLC^AA^, as well as the R163A variant, either to the Golgi complex (Fig.7E and 7F) or to the ER (Fig.7G and 7H) significantly dropped the levels of FAPP1_PH_-GFP at the Golgi complex. These data suggest that PtdIns4P levels at the Golgi complex are sensitive to the local PtdIns content of the Golgi membranes, which is consistent with the relatively low affinity of PI4KB towards PtdIns (Downing et al., 1996; Zhao et al., 2000). Additionally, similar to the effects observed in the outer mitochondrial membrane using the BRET-based measurements, it is interesting that the Golgi-associated localization of both the GFP-*Bc*PI-PLC^H82A^ probe and FAPP1_PH_-GFP were selectively reduced by the recruitment of the inactive FKBP-*Bc*PI-PLC^AA^ H32A directly to the surface of the Golgi, but not to the ER (Fig.7A,7C, 7E, and 7G). This minor effect of the recruited inactive enzyme was attributed to the ability of the recruited FKBP-*Bc*PI-PLC^AA^ H32A enzyme to compete for binding of the *Bc*PI-PLC^H82A^ probe to the resident PtdIns and perhaps masking it from local effectors, including the PI4Ks.

### The steady-state levels of PtdIns are low within the PM

Perhaps the most striking observation using the GFP-*Bc*PI-PLC^H82A^ construct as a PtdIns probe was the lack of significant localization to the PM (Fig.8A). Based on the well-documented importance and relative enrichment of PPIn lipids within the PM, this was unexpected as it has long been assumed from studies of red blood cell membranes (King et al., 1987) that there is a significant reserve pool of PtdIns within the PM. However, recent lipidomics analyses using advanced membrane isolation techniques have already questioned these assumptions by showing that the levels of PtdIns within enriched “PM lawns” were extremely low (Saheki et al., 2016). To further investigate this question, we recruited the FKBP-*Bc*PI-PLC^AA^ R163A to the PM and measured local DAG production. This analysis showed only a minor increase of DAG in the PM that was only detectable using the BRET-based approach (PM-DAG^BRET^, L_10_-mVenus-T2A-sLuc-PKDC1_ab_; Fig.8F). Furthermore, recruitment of a truncated FKBP-PI4KA^ΔN^ to the PM, also failed to alter the PM levels of PtdIns4P (PM-PI4P^BRET^, L_10_-mVenus-T2A-sLuc-P4M_SidM_^x2^; Fig.8B). These data are consistent with a low resting availability of PtdIns within the cytosolic leaflet of the PM. With this conclusion in mind, we sought to monitor the PM content of PtdIns in response to biochemical or pharmacological manipulations of PPIn-modifying enzymes. First, we used an established enzymatic system to rapidly dephosphorylate PPIn species within the PM and simultaneously monitored PtdIns levels. Acute PM recruitment of the engineered enzyme FKBP-Pseudojanin (Hammond et al., 2012), which possesses both 4- and 5-position PPIn phosphatase activities, caused an acute increase of *Bc*PI-PLC^H82A^ levels within the PM (PM-H82A^BRET^, L_10_-mVenus-T2A-sLuc-*Bc*PI-PLC^H82A^) relative to both the baseline and an inactive enzymatic control (FKBP-Pseudojanin^DEAD^; Fig.8C); although the latter still appeared to have some minor effect that could be related to residual 5-phosphatase activity that has been associated with the original Pseudojanin^DEAD^ enzyme scaffold.

**Figure 8.**
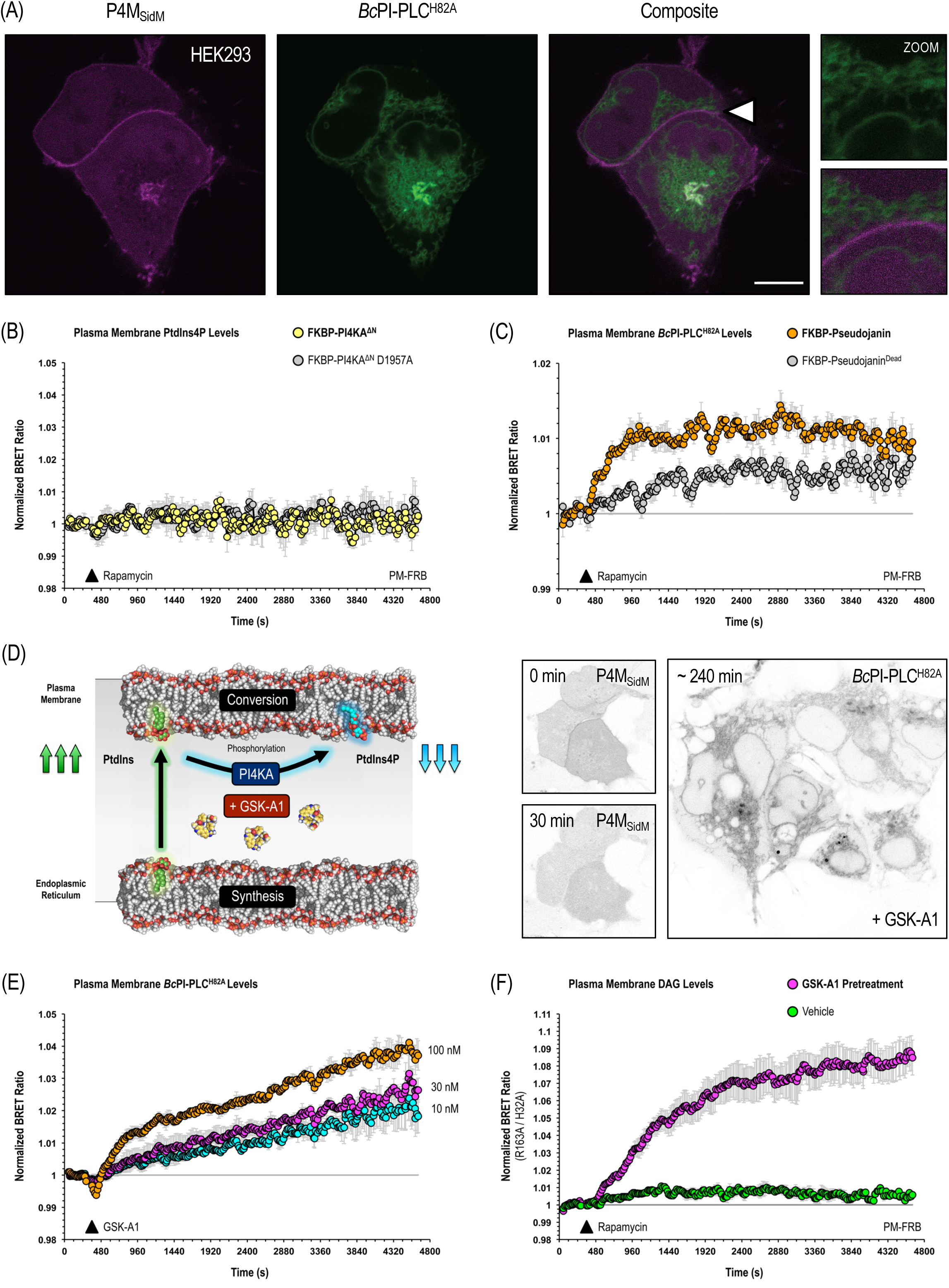
The steady-state levels of PtdIns are low within the PM. (A) Co-expression of EGFP-*Bc*PI-PLC^H82A^ with the unbiased PtdIns4P-binding probe, mCherry-P4M_SidM_, in HEK293-AT1 cells (10 μm scale bar). Despite the co-localization of EGFP-*Bc*PI-PLC^H82A^ with the PtdIns4P pool labelled by mCherry-P4M_SidM_ at the Golgi complex, an enlarged view of the region identified by the white arrowhead clearly shows that the *Bc*PI-PLC^H82A^ probe does not localize to the PM. (B) PtdIns4P production in the PM was measured using the PM-PtdIns4P^BRET^ biosensor (L_10_-mVenus-T2A-sLuc-P4M_SidM_^X2^) together with the iRFP-FKBP-PI4KA^ΔN^ and PM-FRB (PM2-FRB-ECFP^W66A^) heterodimerization system. Consistent with a low resting availability of PtdIns within this compartment, recruitment of FKBP-PI4KA^ΔN^ did not alter the PM levels of PtdIns4P. (C) *Bc*PI-PLC^H82A^ levels within the PM were measured using the PM-H82A^BRET^ biosensor (L_10_-mVenus-T2A-sLuc-*Bc*PI-PLC^H82A^) in combination with the multicistronic L_10_-FRB-T2A-mRFP-FKBP-Pseudojanin heterodimerization system. Relative to both the baseline and the inactive control (L_10_-FRB-T2A-mRFP-FKBP-Pseudojanin^DEAD^), recruitment of FKBP-Pseudojanin, which possesses both 4- and 5-position PPIn phosphatase activities, caused an acute increase of *Bc*PI-PLC^H82A^ levels within the PM. These data are consistent with a localized production of PtdIns following the targeted dephosphorylation of the resident PPIn species, PtdIns4P and PtdIns(4,5)P_2_. (D) Given the need for sustained PPIn production of within the PM, we hypothesized that local delivery of PtdIns would be intimately tied to PI4KA activity and that PtdIns would accumulate within the PM if its rapid conversion to PtdIns4P is prevented. Acute inhibition of PI4KA activity using the selective inhibitor GSK-A1 (100 nM) caused the rapid loss of mCherry-P4M_SidM_ from the PM after 20-30 min and also resulted in the uniform accumulation of EGFP-*Bc*PI-PLC^H82A^ within the PM of HEK293-AT1 cells after roughly 180-240 min. (E) Kinetic analysis using the PM-H82A^BRET^ biosensor (L_10_-mVenus-T2A-sLuc-*Bc*PI-PLC^H82A^) revealed a dose-dependent increase in the PM levels of *Bc*PI-PLC^H82A^ following treatment with GSK-A1. (F) DAG production in the PM was measured using the PM-DAG^BRET^ biosensor (L_10_-mVenus-T2A-sLuc-PKDC1_ab_) in combination with the iRFP-FKBP-*Bc*PI-PLC^AA^ R163A and PM-FRB (PM2-FRB-ECFP^W66A^) heterodimerization system. Consistent with the inability to locally produce PtdIns4P within the PM using the FKBP-PI4KA^ΔN^, recruitment of the FKBP-*Bc*PI-PLC^AA^ R163A failed to elicit a major change in local DAG levels. However, pre-treatment of cells with GSK-A1 (100 nM) for 30 min resulted in a large increase in DAG production within the PM in response to recruitment of the FKBP-*Bc*PI-PLC^AA^ R163A; consistent with an marked increase in PtdIns availability following the inhibition of PtdIns conversion to PtdIns4P by the PM-localized PI4KA. Please note that for each of the BRET experiments shown (B, C, E, and F), drug treatments were added manually after a 4 min baseline BRET measurement; with the post-application measurements beginning at ∼360s and continuing for 60 min (15 s / cycle). All measurements were carried out in triplicate wells (HEK293-AT1; 1×10^5^ cells/well) and repeated in three independent experiments. BRET ratios were first expressed relative to the baseline BRET measurement and then treated wells were normalized to the drug vehicle controls.

Next, we hypothesized that perhaps rapid conversion of PtdIns to PtdIns4P in the PM by PI4KA contributes to the low level of PtdIns that is maintained within the PM (Fig.8D). To test this, we used the selective PI4KA inhibitor, GSK-A1 (Bojjireddy et al., 2014), which gradually drops PM PtdIns4P levels. Treatment with GSK-A1 (100 nM) resulted in a gradual enrichment of *Bc*PI-PLC^H82A^ within the PM that was clearly apparent in almost all cells after 180 to 240 minutes (Fig.8D). To better address the kinetics of these changes, we repeated these experiment using the PM-H82A^BRET^ biosensor and saw a rapid and dose-dependent increase of *Bc*PI-PLC^H82A^ in the PM after acute application of GSK-A1 (Fig.8E). To confirm these findings, we measured DAG production in response to recruitment of FKBP-*Bc*PI-PLC^AA^ R163A after a pre-incubation with GSK-A1. As discussed above, recruitment of the FKBP-*Bc*PI-PLC^AA^ R163A to the PM only evoked a minor increase in the PM level of DAG. In contrast, the same manipulation evoked a rapid and sizeable increase in DAG content within the PM following a 30 min pretreatment of cells with GSK-A1 (100 nM; Fig.8F). These findings strongly support the conclusion that steady-state levels of PtdIns are low within the PM and provide evidence for the PM accumulation of PtdIns in response to the inhibition of PI4KA.

### ER-PM transport of PtdIns is directly channeled towards PPIn production within the PM

Upon activation of selective classes of GPCRs or RTKs, mammalian PLCs cleave the polar head group of PtdIns(4,5)P_2_ to produce two important intracellular second messengers: inositol 1,4,5-trisphosphate (Ins(1,4,5)P_3_) and DAG (Berridge and Irvine, 1984; Berridge, 2016). To maintain the supply of PtdIns(4,5)P_2_ and PtdIns4P within the PM during sustained receptor activation, PtdIns must be delivered and converted to PtdIns4P in the PM (Michell, 1975). The contribution of PtdIns delivery from the ER for the maintenance of PPIn lipids is especially important if the resting level of PtdIns in the PM is indeed low. To determine the impact of acute PtdIns depletion on the turnover of PPIn species, we monitored PtdIns4P (PM-PI4P^BRET^, L_10_-mVenus-T2A-sLuc-P4M_SidM_^x2^; Tóth et al., 2016) or PtdIns(4,5)P_2_ (PM-PI(4,5)P_2_^BRET^, L_10_-mVenus-T2A-sLuc-PLCδ1_PH_; Tóth et al., 2016) levels within the PM after recruitment of FKBP-*Bc*PI-PLC^AA^ R163A to either the PM or indirectly to the ER. Direct recruitment of FKBP-*Bc*PI-PLC^AA^ R163A to the PM failed to alter the PM levels of either PtdIns4P or PtdIns(4,5)P_2_. In contrast, recruitment of FKBP-*Bc*PI-PLC^AA^ R163A to the ER selectively reduced PtdIns4P levels within the PM without altering PtdIns(4,5)P_2_ levels (Fig.9A, 9B, 9C, and 9D). For comparison, treatments with GSK-A1 (100 nM), to inhibit PI4KA, or the activation of endogenous PLC activity using angiotensin-II (AngII; 100 nM) were used to achieve complete loss of PtdIns4P or PtdIns(4,5)P_2_ from the PM, respectively (Fig.9A and 9D). Representative confocal images further support the selective reduction of PtdIns4P by ER recruitment of FKBP-*Bc*PI-PLC^AA^ R163A, as evidenced from the re-localization of the GFP-P4M_SidM_^x2^ probe into the cytosol and towards the substantial PtdIns4P pool at the Golgi complex (Fig.9C). Overall, these data highlight the tight relationship between PtdIns supply from the ER and PtdIns4P production at the PM and are also consistent with previous studies describing the dissociation of PtdIns4P and PtdIns(4,5)P_2_ levels following genetic (Nakatsu et al., 2012; Alvarez-Prats et al., 2018) or pharmacological (Bojjireddy et al., 2014) ablation of PM-associated PI4KA activity. Lastly, we also wanted to examine the PM levels of the *Bc*PI-PLC^H82A^ probe in response to PLC-dependent consumption of PPIn lipids (Balla et al., 1988). Quantified responses, measured using the PM-H82A^BRET^ biosensor, show only minor changes in PM levels of *Bc*PI-PLC^H82A^ compared to the massive relative changes observed in both PtdIns4P and PtdIns(4,5)P_2_ after GPCR-dependent PLC activation (Fig.9G). As shown in more detail with an expanded scale (Fig. 9H), PtdIns initially shows a slight increase, but soon declines slightly below its initial levels within the PM as PPIn re-synthesis begins to take place. Lastly, representative images after stimulation with AngII (100 nM) show that despite a massive increase in the PM levels of DAG, accumulation of *Bc*PI-PLC^H82A^ is not observed within the PM (Fig.9I). This result clearly shows that without PtdIns present within the membrane, acute DAG production or changes to membrane packing, in isolation, are not sufficient to localize the *Bc*PI-PLC^H82A^ probe to the membrane.

**Figure 9.**
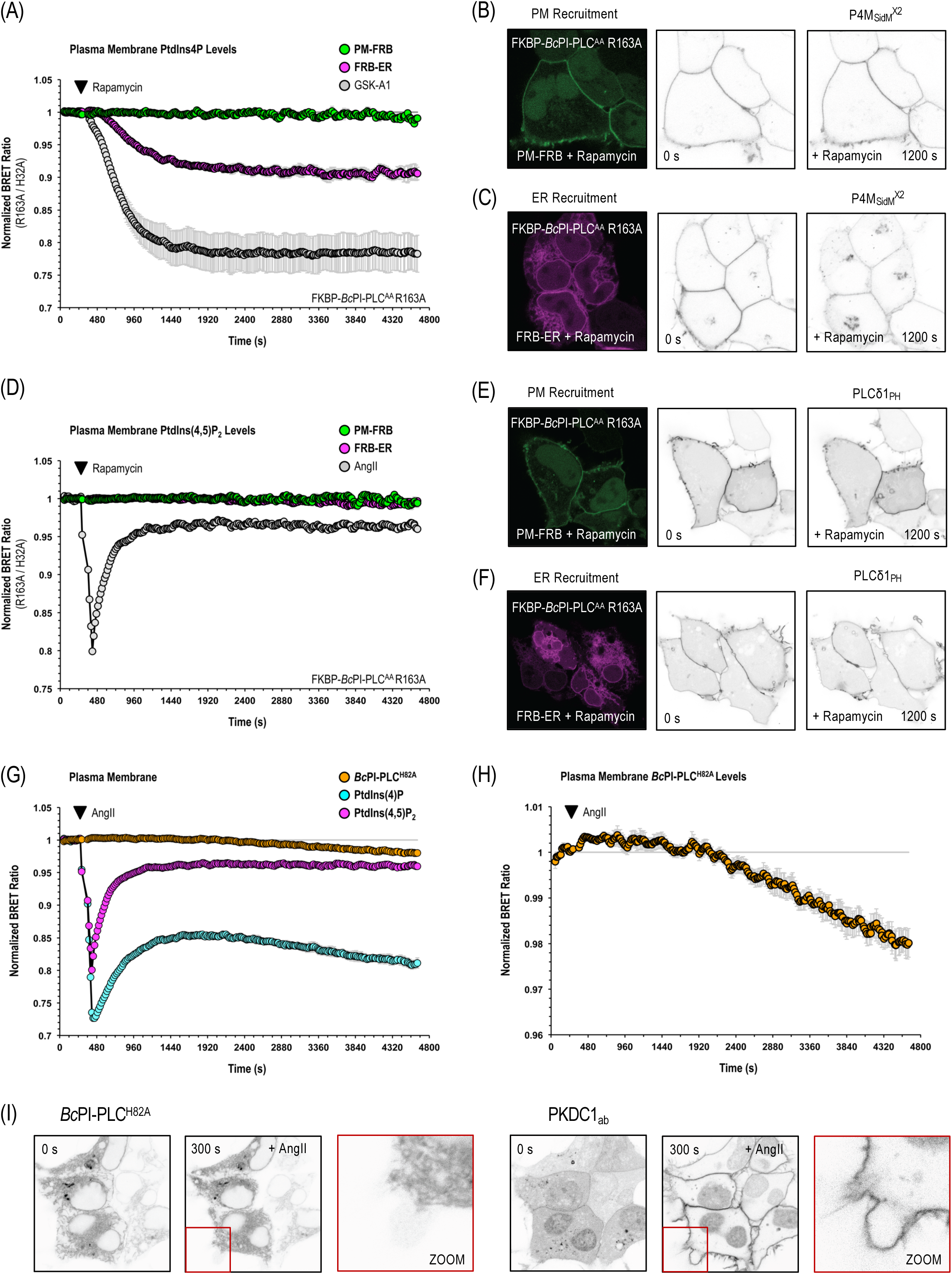
PtdIns delivered to the PM is channeled towards PPIn production. (A) PtdIns4P and (D) PtdIns(4,5)P_2_ levels in the PM were measured using the PM-PtdIns4P^BRET^ (L_10_-mVenus-T2A-sLuc-P4M_SidM_^X2^) or PM-PtdIns(4,5)P_2_^BRET^ (L_10_-mVenus-T2A-sLuc-PLCδ1_PH_) biosensors in response to recruitment of iRFP-FKBP-*Bc*PI-PLC^AA^ R163A either directly to the PM (PM-FRB, PM2-FRB-ECFP^W66A^) or indirectly to the site of PtdIns synthesis within the ER (FRB-ER, mTagBFP2^E215A^-FRB-CyB5_tail_). Treatment with GSK-A1 (100 nM), which selectively inhibits PI4KA, or angiotensin-II (AngII; 100 nM) are included as positive controls for the PM-PtdIns4P^BRET^ and PM-PtdIns(4,5)P_2_^BRET^ biosensors, respectively, as well as to provide scale for any changes to PPIn levels that are associated with the membrane recruitment of FKBP-*Bc*PI-PLC^AA^ R163A. Images of representative cells are shown with the resulting post-rapamycin (100 nM) enzyme recruitment to the PM (right panels; B, PtdIns4P; E, PtdIns(4,5)P_2_) or ER (right panels; C, PtdIns4P; F, PtdIns(4,5)P_2_), as well as the initial (center panels) and final (left panels) localization of the established EGFP-P4M_SidM_^X2^ (B and C) or PLCδ1_PH_-EGFP (E and F) probes following 20 min of localized PtdIns hydrolysis. Although direct recruitment of FKBP-*Bc*PI-PLC^AA^ R163A to the PM did not alter PPIn levels, ER recruitment resulted in the selective depletion of PtdIns4P, but not PtdIns(4,5)P_2_, within the PM. These data suggests that PtdIns delivery is efficiently channeled towards PI4P production and, as shown previously, that PtdIns4P and PtdIns(4,5)P_2_ levels can be uncoupled from one another within the PM. Kinetic analyses of *Bc*PI-PLC^H82A^ (G and H; PM-H82A^BRET^; L_10_-mVenus-T2A-sLuc-*Bc*PI-PLC^H82A^), PtdIns4P (H; PM-PtdIns4P^BRET^; L_10_-mVenus-T2A-sLuc-P4M ^X2^) and PtdIns(4,5)P_2_ (H; PM-PtdIns(4,5)P_2_^BRET^; L_10_-mVenus-T2A-sLuc-PLCδ1_PH_) levels within the PM were measured after treatment with AngII (100 nM). Acute stimulation of receptor-mediated PtdIns(4,5)P_2_ hydrolysis is associated with a minor and transient increase of *Bc*PI-PLC^H82A^ within the PM that is followed by a decrease in total *Bc*PI-PLC^H82A^ levels, which likely reflects the net consumption of PtdIns during sustained receptor activation and the subsequent re-synthesis of PPIn species. (I) Given the role of interfacial activation for membrane binding, representative images showing the localization of the EGFP-*Bc*PI-PLC^H82A^ (left panels) and mRFP-PKDC1_ab_ (right panels) probes in response to stimulation with AngII (100 nM) are shown. Despite the massive increase in the PM levels of DAG (red inset, right), a coordinate accumulation of *Bc*PI-PLC^H82A^ is not observed (red inset, left), which supports an important role for membrane PtdIns content, rather than DAG production or changes to membrane packing in isolation, for the localization of the *Bc*PI-PLC^H82A^ probe. Please note that for each of the BRET experiments shown (A, D, G, and H), drug treatments were added manually after a 4 min baseline BRET measurement; with the post-application measurements beginning at ∼360s and continuing for 60 min (15 s / cycle). All measurements were carried out in triplicate wells (HEK293-AT1; 1×10^5^ cells/well) and repeated in three independent experiments. BRET ratios were first expressed relative to the baseline BRET measurement and then treated wells were normalized to the vehicle-alone controls. Additionally, to facilitate direct comparisons between the (A) PtdIns4P and (D) PtdIns(4,5)P_2_ measurements made in response to either PM or ER recruitment of FKBP-*Bc*PI-PLC^AA^ R163A, values obtained from the recruitment of the active enzyme were also normalized to control measurements made simultaneously using the catalytically-inactive FKBP-*Bc*PI-PLC^AA^ H32A variant.

### Endosomal PtdIns3P levels depend on PtdIns delivery from the ER

Apart from PtdIns4P and PtdIns(4,5)P_2_, monophosphorylated PtdIns3P comprises the largest remaining PPIn pool at roughly 20-30% of the total membrane PtdIns4P content in higher eukaryotes (Sarkes and Rameh, 2010). PtdIns3P is found primarily in early endosomes (Gillooly et a., 2000) and autophagosomes (Funderburk et al., 2010), but the possible presence of PtdIns3P in membranes of the Golgi and ER has also been documented using membrane fractionation (Sarkes and Rameh, 2010). Within the endo-lysosomal system, PtdIns3P is produced primarily within early endosomes by the class III phosphoinositide 3-kinase (Vps34) complex (Volinia et al., 1995) and plays a central role in the spatial control of endosomal fusion and cargo sorting (Schink et al., 2016). In particular, production of PtdIns3P within Rab5-associated endosomes plays a central role in the multistep mechanism that drives endosomal maturation by facilitating the membrane recruitment of protein machineries responsible for triggering the switch from a Rab5-GTP to Rab7-GTP membrane coat (Rink et al., 2005; Poteryaev et al., 2010). Similar to our efforts to understand the role of PtdIns delivery for PPIn synthesis within the Golgi complex and PM, we sought to define the relative contribution of local endosomal PtdIns content and ER-derived PtdIns synthesis for the maintenance of PtdIns3P within Rab5- and Rab7-positive endosomal compartments. Overexpression of wild-type Rab5 (Fig.10A) or Rab7 (Fig.10D) shows minimal co-localization of these endosomal markers with the *Bc*PI-PLC^H82A^ probe; although intimate contacts between the ER and both the early as well as late endosomal compartments are apparent (Fig.10A and 10D, white arrowheads). Accordingly, rapamycin-dependent recruitment of FKBP-*Bc*PI-PLC^AA^ R163A showed a very minor increase in DAG production within Rab5-positive compartments (Rab5-DAG^BRET^, sLuc-PKDC1_ab_-T2A-mVenus-Rab5; Fig.10B). However, a significant increase in DAG levels were seen in Rab7-positive membranes (Rab7-DAG^BRET^, sLuc-PKDC1_ab_-T2A-mVenus-Rab7; Fig.10E). These data are consistent with a a detectable amount of PtdIns within the Rab7-labelled endosomes and could also highlight a role for a continued supply of PtdIns to late endosomes as a result of an increase in the number of contacts with the ER that occurs during endosomal maturation (Friedman et al., 2013; Raiborg et al., 2015). To define the relative roles of PtdIns originating from the surface of endosomes and that transported from the ER for local PtdIns3P production, we monitored PtdIns3P levels within both Rab5- and Rab7-positive endosomes after directly recruiting FKBP-*Bc*PI-PLC^AA^ R163A to the surface of endosomes or indirectly to the ER. Similar to the other monophosphorylated PPIn pools, recruitment of FKBP-*Bc*PI-PLC^AA^ R163A to the ER significantly reduced PtdIns3P content in both the Rab5 (Rab5-PtdIns3P^BRET^, sLuc-FYVE_Hrs_^x2^-T2A-mVenus-Rab5) and Rab7 (Rab7-PtdIns3P^BRET^, sLuc-FYVE_Hrs_^x2^-T2A-mVenus-Rab7) compartments; whereas only minor reductions were observed following the direct recruitment of FKBP-*Bc*PI-PLC^AA^ R163A directly to the respective endosome populations (Fig.10C and 10F). As an important measure of both the selectivity and sensitivity of these BRET biosensors, treatment with a highly selective inhibitor of Vps34 (VPS34-IN1; Bago et al., 2014) significantly reduced the levels of PtdIns3P in both the Rab5- and Rab7-positive endosomes (Fig.10C and 10F). Not surprisingly, the severity of the drop in PtdIns3P content caused by direct inhibition of Vps34 activity exceeded the effects associated with recruitment of the FKBP-*Bc*PI-PLC^AA^ R163A and, based on the relative size of the responses, are consistent with the notion that the PtdIns3P levels are highest within the Rab5-labelled early endosomes. The rapid conversion of PtdIns and the increased PtdIns3P content in the early endosomes might help to explain the relatively low steady-state levels of PtdIns within the Rab5 compartment.

**Figure 10.**
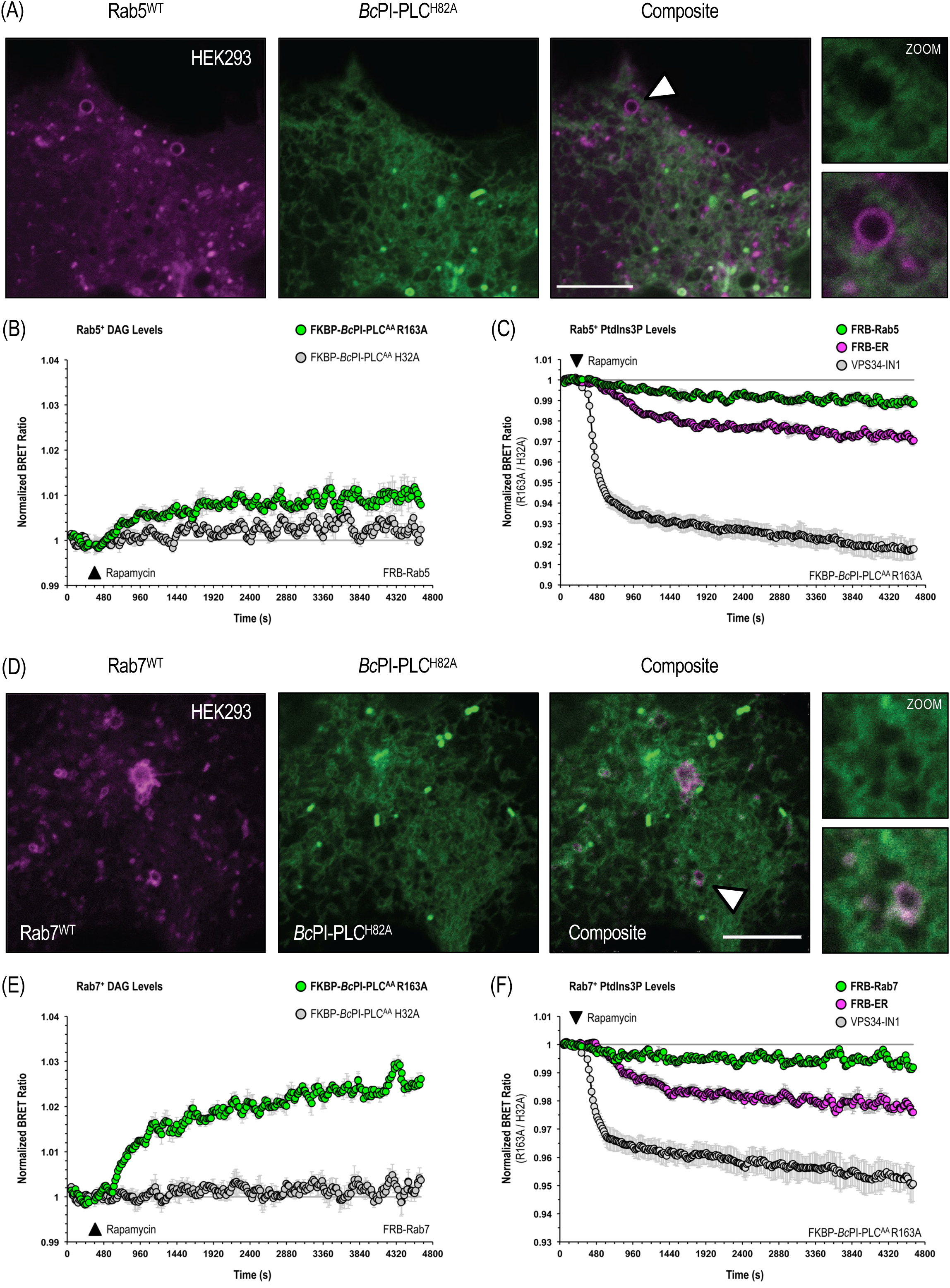
PtdIns3P levels in Rab5- and Rab7-positive endosomes are maintained by the delivery of PtdIns from the ER. Co-expression of the EGFP-*Bc*PI-PLC^H82A^ probe with either (A) mCherry-Rab5^WT^ or (D) mCherry-Rab7^WT^ in HEK293-AT1 cells (5 μm scale bars). Enlarged views of the regions identified by the white arrowheads shows the relative absence of the EGFP-*Bc*PI-PLC^H82A^ probe from Rab5- or Rab7-labelled vesicles; although intimate contacts between Rab-positive compartments and the tubular ER are apparent. DAG production within the (B) Rab5- or (E) Rab7-labelled endosomal compartments were measured using the Rab5-DAG^BRET^ (sLuc-PKDC1_ab_-T2A-mVenus-Rab5) or Rab7-DAG^BRET^ (sLuc-PKDC1_ab_-T2A-mVenus-Rab7) biosensors in combination with iRFP-FKBP-*Bc*PI-PLC^AA^ R163A and either the iRFP-FRB-Rab5 or iRFP-FRB-Rab7 recruiters, respectively. Recruitment of FKBP-*Bc*PI-PLC^AA^ R163A to the Rab5 compartment resulted in only a minor elevation of local DAG levels, whereas recruitment to the Rab7-labelled endosomes caused a more substantial increase in DAG production. Kinetic analyses of PtdIns3P levels within the Rab5 (C; Rab5-PtdIns3P^BRET^; sLuc-FYVE_Hrs_^X2^-T2A-mVenus-Rab5) and Rab7 (F; Rab7-PtdIns3P^BRET^; sLuc-FYVE_Hrs_^X2^-T2A-mVenus-Rab7) compartments are shown in response to the recruitment of iRFP-FKBP-*Bc*PI-PLC^AA^ R163A either directly to the surface of endosomes (C, iRFP-FRB-Rab5; F, iRFP-FRB-Rab7) or indirectly to the site of PtdIns synthesis within the ER (FRB-ER, mTagBFP2^E215A^-FRB-CyB5_tail_). Treatment with the selective class III PtdIns 3-kinase inhibitor VPS34-IN1 (300 nM) is included as both a positive control for the Rab5-PtdIns3P^BRET^ and Rab7-PtdIns3P^BRET^ biosensors, as well as to provide scale for the magnitude of any changes in PtdIns3P levels associated with acute recruitment of FKBP-*Bc*PI-PLC^AA^ R163A. Overall, direct recruitment of FKBP-*Bc*PI-PLC^AA^ R163A to the surface of endosomes resulted in only minor changes to local PtdIns3P levels, while ER recruitment of the active enzyme resulted in a much more substantial decrease in endosomal PtdIns3P contents. Please note that for each of the membrane recruitment studies shown using BRET-based measurements (B, C, E, and F), rapamycin (100 nM) was added manually after a 4 min baseline BRET measurement; with the post-rapamycin measurements beginning at ∼360s and continuing for 60 min (15 s / cycle). All measurements were carried out in triplicate wells (HEK293-AT1; 1×10^5^ cells/well) and repeated in three independent experiments. BRET ratios were first expressed relative to the baseline BRET measurement and then rapamycin-treated wells were normalized to the DMSO-treated controls. Additionally, to facilitate direct comparisons between the (C) Rab5- and (F) Rab7-associated measurements of the PtdIns3P levels made in response to either PM or ER recruitment of the FKBP-*Bc*PI-PLC^AA^ R163A enzyme, values obtained from recruitment of the active enzyme were also normalized to control measurements made simultaneously using the catalytically-inactive FKBP-*Bc*PI-PLC^AA^ H32A variant.

## Discussion

Within mammalian cell membranes, PtdIns represents roughly 10-15 mol% of total phospholipids (Vance, 2015), while the diverse PPIn species only comprise an estimated 2-5% of the available PtdIns (Xu et al., 2003; Sarkes and Rameh, 2010). The relative abundance of PtdIns and its role as a precursor for PPIn production, has led to the assumption that PtdIns is fairly abundant in membranes where the various PPIn lipids exist. Early pioneering studies to visualize the subcellular distribution of PtdIns involved the synthesis of a fluorescent PtdIns analog and revealed accumulation of the tagged lipid within membranes corresponding to the ER, mitochondria, and perinuclear Golgi region (Uster and Pagano, 1986; Ting and Pagano, 1990). However, an examination of total lipid contents after the exogenous delivery of the labelled PtdIns suggested that conversion of the fluorescently-tagged PtdIns to DAG was required for observing the labeling of intracellular membranes (Ting and Pagano, 1990). The apparent hydrolysis of the labelled PtdIns was most prominent in Swiss 3T3 cells and led to the hypothesis that a cell-surface or secreted PI-PLC could be responsible for remodeling the exogenous PtdIns (Ting and Pagano, 1990). These unresolved discrepancies, as well as the need to incorporate the fluorescent label into the fatty acid side chains, question the reliability of these approaches for mapping the intracellular distribution of PtdIns. As a result, we looked for an alternative method for the visualization of intracellular PtdIns and eventually devised a protein-engineering platform that utilizes the substrate selectivity of the bacterial *Bc*PI-PLC to complete comprehensive investigations of PtdIns distribution and availability in live cells.

Efforts to visualize membrane-embedded PtdIns involved the rational design of a *Bc*PI-PLC variant, *Bc*PI-PLC^H82A^, possessing a mutation within the active site that successfully removes catalytic activity without interfering with the coordination of the lipid substrate, PtdIns (Fig.S1). It is important to note that with this approach, the localization of the probe reports on the PtdIns abundance only at the cytoplasmic leaflet of organelle membranes. Our results using the engineered *Bc*PI-PLC^H82A^ probe suggest that the resting levels of PtdIns within the PM and endosomes are kept extremely low and, in addition to the site of PtdIns synthesis within the ER, we identify a relative enrichment of PtdIns in the membranes of the Golgi complex, peroxisomes, and mitochondria. This pattern of the PtdIns distributions is in general agreement with membrane fractionation studies that also define the ER and Golgi complex as containing the highest percentages of PtdIns (∼9%) among organelle compartments; exceeding those associated with the endolysosomal system, nucleus, and PM (∼4-7%; Vance, 2015). It is important to mention that these fractionation values refer to the sum of outer and inner leaflets of the respective membranes. However, while bulk measurements show PtdIns as a relatively minor component of mitochondrial membranes, estimated at ∼5-7% of the total lipid contents, isolation of purified inner and outer membranes reveal an enrichment of PtdIns in the outer mitochondrial membrane (∼9-13% total phospholipids) relative to the inner membrane (∼2-5% total phospholipids; Ardail et al., 1990; de Kroon et al., 1997; Daum and Vance, 1997). Based on our current study, within the cytosolic leaflet, the mitochondria appear to possess a relative PtdIns content similar to that associated with bulk membranes of the ER and Golgi complex. Our findings that, when compared with the ER membrane, PtdIns concentrations appear to be higher within the cytosolic leaflet of membranes forming the Golgi complex, peroxisomes, and mitochondria raises the questions of how these enrichments of PtdIns occur and whether these PtdIns sources serve an unrecognized function. Although the Golgi has a high PtdIns4P content that undergoes rapid turnover, this alone may not explain the high PtdIns levels, since the PM and endosomes also have high PPIn levels and yet, limited amounts of PtdIns were detected in these membrane compartments. Interestingly, a series of studies have shown the important regulatory role of DAG in the control of Golgi functions (Baron and Malhotra, 2002; Bossard et al., 2007; Fernández-Ulibarri et al., 2007; Asp et al., 2009) and it has been suggested that activation of PLC enzymes, which are sensitive to Gβγ subunits, generate DAG from PPIn species at the Golgi (Díaz Añel, 2007). Some of these studies also claim that the substrate used by the Golgi-resident PLC isoforms is PtdIns4P (Zhang et al., 2013; Sicart et al., 2015), but direct hydrolysis of PtdIns by some of the mammalian PLCs cannot be ruled out.

The physiological functions of PtdIns in the mitochondria and peroxisomes are even more intriguing given that enzymes directly using PtdIns as a substrate have not been described to function within, or interact with, the outer membrane of mitochondria. It is notable, though, that multiple proteomic screens have identified PI4K isoforms within mitochondria-enriched datasets (Schon, 2007; Calvo et al., 2016; Hung et al., 2017). Downstream of PtdIns, specific isoforms or alternatively-spliced variants of the PPIn phosphatases synaptojanin 2 (Synj2A; Nemoto and De Camilli, 1999) and phosphatase and tensin homolog deleted on chromosome 10 (PTEN and PTENα; Bononi et al., 2013; Liang et al., 2014) have been shown to localize to the mitochondria, but, apart from scaffolding or allosteric functions, the explicit roles for their lipid phosphatase activities remains largely uncharacterized. The localization of these enzymes is also interesting given that, using the available probes, no PPIn species has ever been detected on the outer surface of mitochondria and our efforts to find PPIn lipids associated with the outer mitochondrial membrane have been unsuccessful. One functional study examining a role for PPIn lipids within the mitochondria provided indirect evidence for PtdIns(4,5)P_2_ as a local regulator of mitochondrial dynamics (Rosivatz and Woscholski, 2011). However, those studies relied on the ectopic overexpression of a mitochondrial-targeted PLCδ1_PH_ domain, which could also function by altering inter-organelle contacts through protein-protein interactions or through localized buffering of Ins(1,4,5)P_3_ levels to change ER-mitochondria associated Ca^2+^ dynamics. Similar to the mitochondria, to date, few studies have looked at the content or functions of PtdIns and PPIn derivatives in peroxisomes. Briefly, fractionation studies show that PtdIns represents ∼5% of the total phospholipid content of the peroxisomes (Hardeman et al., 1990) as well as suggest, relative to other inositol-containing lipids, that the levels of PtdIns4P, PtdIns(3,5)P_2_, and PtdIns(4,5)P_2_ are enriched in peroxisomal membranes (Jeynov et al., 2006). The potential for a specific enrichment of PPIn lipids is supported by recent reports describing the interaction of PtdIns(4,5)P_2_ present on the peroxisome surface with the lysosomal protein synaptotagmin VII to establish contact sites between lysosomes and peroxisomes (Chu et al., 2015; Hu et al., 2018). More studies will be needed to define the role of PtdIns and PPIn species in these compartments.

Considering that the biosynthetic machinery required for PtdIns production is almost exclusively localized to the ER, the low abundance of PtdIns in membranes where PPIn species play critical regulatory roles, such as in the PM or endosomes raises the question whether PtdIns delivery to these compartments is part of the process by which PPIn generation is controlled. Our previous studies identifying a mobile ER-derived PIS compartment that would be an ideal candidate to serve as a PtdIns delivery platforms (Kim et al., 2011). It is important to note that no PtdIns was detected using the *Bc*PI-PLC^H82A^ probe in the dynamic PIS-positive structures. This finding may suggest that PtdIns is rapidly transferred from this active compartment to acceptor membranes where it is again quickly converted to either PtdIns4P, such as in the PM or Golgi complex, or PtdIns3P in early endosomes. It is also noteworthy that, except for the Golgi complex, PtdIns appears to be enriched in membranes that lack significant local pools of PPIn lipids. Therefore, it is possible that the relatively low PtdIns content associated with the PM and endosomal system reflects the fact that local delivery of PtdIns is tightly coupled to the production of phosphorylated PPIn derivatives. In this context, it is worth reiterating the fact that it was more effective to alter the levels of PtdIns4P in the PM and PtdIns3P levels in endosomes by consuming PtdIns at the ER rather than trying to intercept PtdIns within the specific membranes where the PPIn lipids are made. These data suggest a direct channeling of the PtdIns substrate to the respective PPIn-producing kinases in the membranes receiving PtdIns and also stresses the importance of understanding how newly synthesized PtdIns produced in the ER reaches other membranes. Vesicular trafficking is an obvious route for PtdIns delivery, especially for communication between the ER and Golgi complex, but non-vesicular lipid transfer by specialized lipid-transfer proteins (LTPs) is increasingly recognized as a plausible alternative, if not the major lipid transport mechanism (Kim et al., 2013; Wong et al., 2019). In fact, the complex interplay between the vesicular and non-vesicular transport systems that converge at the Golgi complex might help to explain the unique enrichment of both PtdIns and PtdIns4P that occurs in this compartment and could also be an indication of functional heterogeneity in the PtdIns pools that exist throughout the ER and Golgi interface. More generally, it seems as though non-vesicular transport of PtdIns is likely to drive the local delivery of PtdIns in diverse membrane contexts. Interestingly, members of the StARkin-related and Sec14 domain-containing (CRAL-TRIO) superfamilies of LTPs have been classically defined as PtdIns-transfer proteins (PITPs) that are thought to function by redistributing PtdIns from donor to acceptor membranes, likely as part of a heterotypic exchange involving additional lipid species such as phosphatidylcholine or phosphatidic acid (Cockcroft and Carvou, 2007; Grabon et al., 2019). Despite their name, it is yet to be understood how these PITPs function within intact cells; although unique PITPs are increasingly linked to specific signaling modalities in studies of model organisms (Xie et al., 2018; Huang et al., 2018). In addition to the autonomous PITPs, recent work has also suggested that PtdIns can be used as a lipid cargo by the relatively promiscuous TUbular LIPid-binding (TULIP) family of LTPs that possess synaptotagmin-like mitochondrial lipid-binding protein (SMP) domains; including a report of preferential PtdIns transfer from the ER to the PM by the SMP domain-containing protein TMEM24 following PLC activation and PtdIns(4,5)P_2_ hydrolysis (Lees et al., 2017). Understanding the cargo selectivity and potential interplay between the diverse families of PtdIns-binding LTPs represents a major goal for future studies. The inability of the FKBP-*Bc*PI-PLC^AA^ to intercept and hydrolyze the delivered PtdIns within the acceptor membrane are not at odds with the proposed model of substrate-presentation by various families of LTPs to membrane associated effectors, including both major superfamilies of PITPs (Panaretou et al., 1997; Bankaitis et al., 2010; Grabon et al., 2015). The ability to directly channel the delivered PtdIns to PPIn-producing or PtdIns-modifying enzymes is tantalizing, however there is not yet a clear structural basis for how the bound PtdIns cargo could be efficiently moved into or interfaced with known PtdIns kinases. While direct presentation of the substrate to effectors is certainly not a necessity, our observations demonstrate the intimate relationship between PtdIns transport from the ER and PPIn turnover using intact cell membranes. These details clearly merit further investigation and could have important translational consequences for defining novel regulatory mechanisms that facilitate the localized production of clinically-relevant PPIn lipids.

Lastly, while our studies suggest that PtdIns is the primary localization signal responsible for recruiting the inactivated *Bc*PI-PLC^H82A^ probe to membrane compartments, it is important to note that PtdIns binding may not be the sole determinant responsible for the membrane affinity of this probe. In particular, *in vitro* studies have shown that the *K_d_* associated with binding of the *Bacillus* PI-PLC to substrate-poor vesicles is significantly lower than the apparent *K_m_* for the hydrolysis of membrane-embedded PtdIns (Qian et al., 1998). These data highlight the fact that membrane recognition by the *Bc*PI-PLC requires a combination of interfacial as well as active site binding. Consequently, at present, it cannot be ruled out that, in addition to the presence of PtdIns, other membrane features, such as DAG content and local membrane packing defects (Lehto and Sharom, 2002; Ahyayauch et al., 2015) or the relative phosphatidylcholine abundance (Zhang et al., 2004; Pu et al., 2009; Yang et al., 2015), may also play a role in the association of the *Bc*PI-PLC^H82A^ probe with membranes. It is important to emphasize that our conclusions regarding the membrane distribution of PtdIns are based on a combination of approaches and not solely on *Bc*PI-PLC^H82A^ localization; even though this probe appears to faithfully map the cellular PtdIns landscape. However, we would still recommend caution when interpreting the results of experiments based solely on the behavior of the *Bc*PI-PLC^H82A^ probe.

## Summary

In this study we present a unique platform for completing molecular investigations of PtdIns metabolism and trafficking that utilizes the substrate selectivity of the *Bc*PI-PLC. Our experiments showed very limited amounts of PtdIns in the PM and endosomes as well as identified PtdIns within the membranes of the ER and an enrichment in the Golgi complex, peroxisomes, and outer mitochondrial membrane. Our results defining the subcellular distribution of PtdIns will pave the way for future studies that will address the functional role of PtdIns within these membrane compartments, including those independent of PPIn production, which may significantly influence local membrane functions or cellular metabolism. In addition to defining the steady-state subcellular pools of PtdIns, we also provide a powerful new strategy to dissect the complex trafficking events that distribute PtdIns between membranes by selectively hydrolyzing distinct intracellular pools of PtdIns. Results collected using these unique molecular approaches have allowed us to examine the relationship between localized PtdIns availability and PPIn turnover. In the future, we hope that our findings defining the distribution and dynamics of PtdIns metabolism, as well as the establishment of these novel molecular tools, will allow for the identification of new regulatory pathways that integrate PtdIns production with more global changes in cellular lipid homeostasis. Additionally, it has not escaped our attention that establishing this organelle-targeted FKBP-*Bc*PI-PLC^AA^ recruitment system provides a unique opportunity to exploit the biophysical properties related to changing the relative abundance of PtdIns and DAG to examine the effects of selectively altering lipid composition on the local control of membrane dynamics. Overall, we hope these significant contributions to PtdIns biology not only enhance our understanding of the lipid landscape of eukaryotic membranes but also serve to inform our conceptions of the molecular mechanisms that control the onset of PPIn-dependent pathologies, as well as other phospholipid-related disorders, which manifest within specific membrane contexts.

## Materials and Methods

### Cell Culture

COS-7 (CRL-1651; ATCC) or HEK293-AT1 cells, which stably express the rat AT_1a_ angiotensin II receptor (Hunyady et al., 2002), were cultured in Dulbecco’s Modified Eagle Medium (DMEM-high glucose; Life Technologies) containing 10% (vol/vol) FBS and supplemented with a 1% solution of penicillin/streptomycin (Gibco, Life Technologies). Alternatively, human HT-1080 fibrosarcoma cells (CCL-121; ATCC) were maintained using Eagle’s Minimum Essential Medium (EMEM; Millipore Sigma) supplemented with 2 mM _L_-glutamine and containing 10% (vol/vol) FBS as well as a 1% solution of penicillin/streptomycin. Each of these cell lines were maintained at 37°C and 5% CO_2_ in a humidified atmosphere. Cell lines are also regularly tested for *Mycoplasma* contamination using a commercially-available detection kit (InvivoGen) and, after thawing, all cell cultures are treated with plasmocin (InvivoGen) at 500 µg/ml for the initial three passages (6-9 days) as well as supplemented with 5 µg/ml of the prophylactic for all subsequent passages.

### Reagents

All compounds were prepared in the indicated solvent and stored in small aliquots at -20°C. Rapamycin (Sigma Millipore; 100 μM stock) and VPS34-IN1 (Selleck Chemicals; 300 μM stock) were dissolved in DMSO. Production and validation of the PI4KA-selective inhibitor, GSK-A1, has been described previously (Bojjireddy et al., 2014) and stock solutions were prepared at 100 μM in DMSO. Coelenterazine h (Regis Technologies) was dissolved in 100% ethanol (vol/vol) at 5 mM. Angiotensin II (Human octapeptide; Bachem) was first dissolved in ethanol at 1mM before being prepared as 100 μM aliquots for storage by dilution with ddH_2_O water. MitoTracker Red (ThermoFisher Scientific) was pre-diluted 1:100 in DMSO from the concentrated stock for storage in small aliquots at -20°C. Diluted solutions of MitoTracker Red were added directly to the medium of transfected cells at a 1:1000 dilution (1:100,000 final concentration) and allowed to equilibrate for 15-30 min at 37°C prior to imaging.

### DNA Constructs

In general, plasmids were constructed by standard restriction cloning using enzymes from New England Biolabs, while site directed mutagenesis was done using the QuickChange II kit (Agilent). Complex reconfigurations of vector backbones and all point mutations were subsequently verified using standard Sanger sequencing (Macrogen, USA). Briefly, the design of the following plasmids have been described elsewhere: EGFP-FAPP1_PH_ (Balla et al., 2005), PM2-FRB-ECFP (Várnai et al., 2006), mRFP-FKBP-5-ptase-dom (Várnai et al., 2006), mRFP-ER(Sac1_1-38_) (Várnai et al., 2007), AKAP-FRB-ECFP (Csordás et al., 2010), mRFP-PI-PLC (*Listeria monocytogenes*; Kim et al., 2011), EGFP-PKDC1_ab_ (Kim et al., 2011), EGFP-P4M_SidM_ (Hammond et al., 2014), mCherry-P4M_SidM_ (Hammond et al., 2014), iRFP-P4M_SidM_ (Hammond et al., 2014), EGFP-P4M_SidM_^X2^ (Hammond et al., 2014), iRFP-FRB-Rab5 (Hammond et al., 2014), iRFP-FRB-Rab7 (Hammond et al., 2014), mCherry-FKBP-PI4KA^ΔN^ (Hammond et al., 2014), mCherry-FKBP-PI4KA^ΔN^ D1957A (Hammond et al., 2014), FRB-mCherry-Giantin_tail_ (Hammond et al., 2014), NES-mdsRed-Spo20^DM^ (Kim et al., 2015), L_10_-mVenus-T2A-sLuc-D4H (Sohn et al., 2018), sLuc-P4M_SidM_^X2^-T2A-mVenus-Rab5 (Baba et al., 2019), sLuc-P4M_SidM_^X2^-T2A-mVenus-Rab7 (Baba et al., 2019), sLuc-FYVE^EEA1^-T2A-mVenus-Rab5 (Baba et al., 2019), and sLuc-FYVE^EEA1^-T2A-mVenus-Rab7 (Baba et al., 2019). Also, we would like to thank the laboratories of Harald Stenmark (EGFP-FYVE_Hrs_^X2^; Gillooly et al., 2000), Jennifer Lippincott-Schwartz (mRFP-SKL; Kim et al., 2006), Takanari Inoue (ECFP-FRB-Giantin; Komatsu et al., 2010), Robert Lodge (mCherry-Rab5^WT^ and mCherry-Rab7^WT^; Hammond et al., 2014), Bianxiao Cui (Addgene plasmid #102250; Duan et al., 2015), Peter Várnai (PLCδ1_PH_-mVenus, L_10_-mVenus-T2A-sLuc-P4M_SidM_^X2^, L_10_-mVenus-T2A-sLuc-PLCδ1_PH_, L_10_-FRB-T2A-mRFP-FKBP-Pseudojanin, and L_10_-FRB-T2A-mRFP-FKBP-Pseudojanin^DEAD^; Tóth et al., 2016), and Gerry Hammond (mTagBFP2-FKBP-CyB5_tail_; Zewe et al., 2018) for generously providing constructs. Alternatively, the cloning procedures used for generating DNA constructs unique to this study are provided below and the primers required for both PCR-mediated cloning or site-directed mutagenesis are listed in Supplemental Table S4.

To facilitate the diverse cloning strategies needed for this study, the *Bacillus cereus (Bc*)PI-PLC (signal peptide removed, residues 32-329; GenBank Accession: AAA22665.1) was codon-optimized for mammalian expression and custom synthesized in the pUCminusMCS vector shuttle by Blue Heron Biotech LLC (Bothell, WA, USA). EGFP-*Bc*PI-PLC was generated by inserting the *Bc*PI-PLC sequence into the pEGFP-C1 vector (Clonetech) using the flanking EcoRI and BamHI restriction sites included in the synthesized pUCminusMCS-*Bc*PI-PLC construct. Site directed mutagenesis of the parent EGFP-*Bc*PI-PLC backbone was done to introduce alanine substitutions at His32 (H32A), His82 (H82A), Trp47 (W47A), and Trp242A (W242A); as well as sequential mutagenesis of both interfacial tryptophan residues to generate EGFP-*Bc*PI-PLC W47A/W242A (^AA^). To facilitate co-localization studies with the various EGFP-*Bc*PI-PLC constructs, mRFP-PKDC1_ab_ and AKAP-mRFP were made using the original cloning sites to digest the respective GFP-tagged variants and insert them into the empty pmRFP-C1 or pmRFP-N1 vectors (Clonetech). Alternatively, the recruitable mRFP- FKBP-*Bc*PI-PLC^AA^ enzyme was created by replacing the Type IV 5-phosphatase domain in the mRFP-FKBP- 5-ptase-dom construct with the *Bc*PI-PLC^AA^ insert amplified from EGFP-*Bc*PI-PLC^AA^ using SacI and KpnI restriction sites. To alleviate any concerns related to potential interference with BRET measurements, we also made iRFP-FKBP-*Bc*PI-PLC^AA^ by replacing the mRFP with the iRFP module from iRFP-P4M_SidM_ using NheI and BspEI restriction sites. Site directed mutagenesis of the iRFP-FKBP-*Bc*PI-PLC^AA^ backbone was used to generate the H32A inactive control as well as the R163A, R163K, Y200A, and Y200F active site mutants for screening. Similarly, for BRET measurements using recruitment of the FKBP-PI4KA, iRFP-FKBP-PI4KA^ΔN^ was generated from mCherry-FKBP-PI4KA^ΔN^ by replacing mCherry with the iRFP insert from iRFP-FKBP-*Bc*PI-PLC^AA^ using AgeI and HindIII restriction sites. Using the resulting iRFP-FKBP-PI4KA^ΔN^ construct, we also made a catalytically inactive mutant by replacing the PI4KA^ΔN^ with the PI4KA^ΔN^ D1957A insert from mCherry-FKBP-PI4KA^ΔN^ D1957A using HindIII and KpnI restriction sites. For the recruitment of FKBP-tagged enzymes, the ER-targeted mTagBFP2-FRB-CyB5_tail_ construct was created by amplifying the FRB insert from AKAP-FRB-ECFP and replacing the FKBP12 module from mTagBFP2-FKBP-CyB5_tail_ using XhoI and EcoRI restriction sites. Peroxisomal targeting was done by creating the PEX-FRB-ECFP recruiter by amplifying the transmembrane peroxisome-targeting domain of PEX3 (residues 1-42; Duan et al., 2015) from PEX-mCherry-CRY2 and inserting this in place of the AKAP sequence in AKAP-FRB-ECFP using NheI and BamHI restriction sites. Mutagenesis of the ECFP (W66A) or mTagBFP2 (E215A) chromophores through alanine substitution was done to inactivate the fluorescence of the AKAP-FRB-ECFP, PEX-FRB-ECFP, ECFP-FRB-Giantin, PM2-FRB-ECFP, and mTagBFP2-FRB-CyB5_tail_ membrane recruitment constructs to prevent interference with the BRET measurements.

The design of the organelle-specific BRET biosensors required several unique cloning strategies. To construct the mito-DAG^BRET^ (AKAP-mVenus-T2A-sLuc-PKDC1_ab_), mito-PtdIns4P^BRET^ (AKAP-mVenus-T2A-sLuc-P4M_SidM_^X2^), and mito-H82A^BRET^ (AKAP-mVenus-T2A-sLuc-*Bc*PI-PLC^H82A^) biosensors, we first generated AKAP-mVenus by exchanging the PLCδ1_PH_ domain from PLCδ1_PH_-mVenus with the mitochondrial-targeting sequence (N-terminal residues 34-63 of mAKAP1; Csordás et al., 2010) from AKAP-mRFP using AgeI and NotI restriction sites. Specifically for the mito-DAG^BRET^ construct, we also needed to make an L_10_-mVenus-T2A-sLuc-PKDC1_ab_ intermediate by replacing the P4M_SidM_^X2^ module of L_10_-mVenus-T2A-sLuc-P4M_SidM_^X2^ with the PKDC1_ab_ domain from EGFP-PKDC1_ab_ using BglII and BamHI restriction sites. We then used an NheI and BsrGI double-digest to insert the AKAP-mVenus fragment in place of the L_10_-mVenus targeting sequence in both L_10_-mVenus-T2A-sLuc-PKDC1_ab_ and L_10_-mVenus-T2A-sLuc-P4M_SidM_^X2^ to make the final AKAP-mVenus-T2A-sLuc-P4M_SidM_^X2^ and AKAP-mVenus-T2A-sLuc-PKDC1_ab_ constructs. Alternatively, AKAP-mVenus-T2A-sLuc-*Bc*PI-PLC^H82A^ was generated by amplifying out the *Bc*PI-PLC^H82A^ insert from EGFP-*Bc*PI-PLC^H82A^ and inserting it in place of the PKDC1_ab_ module from AKAP-mVenus-T2A-sLuc-PKDC1_ab_ using SpeI and BamHI restriction sites. Production of the ER-DAG^BRET^ (sLuc-PKDC1_ab_-T2A-mVenus-CyB5_tail_), Golgi-DAG^BRET^ (sLuc-PKDC1_ab_-T2A-mVenus-Giantin), and PEX-DAG^BRET^ (PEX3_1-42_-mVenus-T2A-sLuc-PKDC1_ab_) biosensors also required sequential steps. First, the CyB5_tail_ or Giantin membrane-targeting fragments were amplified from mTagBFP2-FKBP-CyB5_tail_ and ECFP-FRB-Giantin, respectively, and inserted in place of Rab7 in sLuc-FYVE^EEA1^-T2A-mVenus-Rab7 using BspEI and KpnI restriction sites. The FYVE^EEA1^domain was then replaced with PKDC1_ab_ in both the sLuc-FYVE^EEA1^-T2A-mVenus-CyB5_tail_ and sLuc-FYVE^EEA1^-T2A-mVenus-Giantin constructs by amplifying the PKDC1_ab_ insert from AKAP-mVenus-T2A-sLuc-PKDC1_ab_ and using PvuI and SalI restriction sites. Alternatively, we utilized a unique PvuI site present in the L_10_-mVenus-T2A-sLuc-Spo20^(DM)^ construct to replace the PM-targeting sequence with the transmembrane PEX3_1-42_ fragment amplified from PEX-mCherry-CRY2 using NdeI and PvuI restriction sites. From the PEX3_1-42_-mVenus-T2A-sLuc-Spo20^(DM)^ intermediate, we replaced the PtdOH-binding Spo20^51-90^ C54S / C82S (Spo20^DM^) cassette with the PKDC1_ab_ domain from AKAP-mVenus-T2A-sLuc-PKDC1_ab_ using BsrGI and BamHI restriction sites. Although not directly used in this study, the L_10_-mVenus-T2A-sLuc-Spo20^(DM)^ construct was designed by amplifying the Spo20^DM^ insert from NES-mdsRed-Spo20^DM^ and replacing the P4M_SidM_^X2^ module in L_10_-mVenus-T2A-sLuc-P4M_SidM_^X2^ using BglII and EcoRI restriction sites. This construct was chosen for mutagenesis to introduce a unique PvuI site between the membrane-targeting and mVenus modules due to the relative abundance of unique cloning sites surrounding the Spo20^DM^ cassette at the N-terminus of this construct. To measure potential changes in the PM localization of the PtdIns-sensitive *Bc*PI-PLC^H82A^ probe, we generated the PM-H82A^BRET^ (L_10_-mVenus-T2A-sLuc-*Bc*PI-PLC^H82A^) biosensor using XhoI and EcoRI restriction sites to replace the cholesterol-binding D4H domain in the L_10_-mVenus-T2A-sLuc-D4H construct with the *Bc*PI-PLC^H82A^ insert amplified from EGFP-*Bc*PI-PLC^H82A^. Within the endosomal system, the Rab5-DAG^BRET^ (sLuc-PKDC1_ab_-T2A-mVenus-Rab5) biosensor was made using PvuI and AgeI restriction sites to swap out the FYVE^EEA1^ domain in sLuc-FYVE^EEA1^-T2A-mVenus-Rab5 for the PKDC1_ab_ domain from sLuc-PKDC1_ab_-T2A-mVenus-CyB5_tail_. Alternatively, for creating the Rab7-DAG^BRET^ (sLuc-PKDC1_ab_-T2A-mVenus-Rab7) biosensor, the FYVE^EEA1^ domain from sLuc-FYVE^EEA1^-T2A-mVenus-Rab7 was exchanged for PKDC1_ab_ from sLuc-PKDC1_ab_-T2A-mVenus-CyB5_tail_ using PvuI and SalI restriction sites. The high-affinity Rab7-PtdIns3P^BRET^ (sLuc-FYVE_Hrs_^X2^-T2A-mVenus-Rab7) biosensor was constructed by replacing the FYVE^EEA1^ module within sLuc-FYVE^EEA1^-T2A-mVenus-Rab7 with the FYVE_Hrs_^X2^ insert amplified from EGFP-FYVE_Hrs_^X2^ using PvuI and SalI restriction sites. Alternatively, to make the high-affinity Rab5-PtdIns3P^BRET^ (sLuc-FYVE_Hrs_^X2^-T2A-mVenus-Rab5) biosensor, the FYVE_Hrs_^X2^-T2A-mVenus insert from sLuc-FYVE_Hrs_^X2^-T2A-mVenus-Rab7 was inserted in place of FYVE^EEA1^-T2A-mVenus in sLuc-FYVE^EEA1^-T2A-mVenus-Rab5 using PvuI and HindIII restriction sites.

### Live-Cell Confocal Microscopy

For imaging, HEK293-AT1 cells (3×10^5^ cells/dish) were plated with a final volume of 1.5 mL on 29 mm circular glass-bottom culture dishes (#1.5; Cellvis) pre-coated with 0.01% poly-L-lysine solution (Sigma), while both the COS-7 (1×10^5^ cells/dish) and HT-1080 (1.5×10^5^ cells/dish) cell lines were plated without any additional coating of the culture dishes. The cells were allowed to attach overnight prior to transfection of plasmid DNAs (0.1-0.2 μg/well) using Lipofectamine 2000 (2-5 μL/well; Invitrogen) within a small volume of Opti-MEM (200 μL; Invitrogen) according to the manufacturer’s instructions, with the slight modification of removing the media containing the Lipofectamine-complexed DNA at 4-6 h post-transfection and replacing it with complete culture medium. In general, cell densities were always kept between 50-80% confluence for the day of imaging. Also, please note that studies using the rapamycin-inducible protein heterodimerization system used a 1:2:1 ratio of plasmid DNA for transfection of the FKBP-tagged enzyme (0.1 μg), FRB-labelled recruiter (0.2 μg), and indicated protein or biosensor of interest (0.1 μg; total DNA: 0.4 μg/well). After 18-20 h of transfection, cells were incubated in 1 mL of modified Krebs-Ringer solution (containing 120 mM NaCl, 4.7 mM KCl, 2 mM CaCl_2_, 0.7 mM MgSO_4_, 10 mM glucose, 10 mM HEPES, and adjusted to pH 7.4) and images were acquired at room temperature using either the Zeiss LSM 710 (63x/1.40 N.A. Plan-Apochromat Oil objective) or Zeiss LSM 880 (63x/1.40 N.A. Plan-Apochromat Oil DIC M27 objective) laser-scanning confocal microscopes (Carl Zeiss Microscopy). Image acquisition was performed using the ZEN software system (Carl Zeiss Microscopy), while image preparation and analysis was done using the open-source FIJI platform (Schindelin et al., 2012). Please note, unless explicitly labeled otherwise, all representative images shown are of HEK293-AT1 cells.

### Measurements of Membrane Lipid Composition using BRET-based biosensors within Intact Cells

The construction of the BRET-based lipid reporters has been detailed above. Briefly, the design of these biosensors allows for the quantitative description of membrane lipid composition within defined subcellular compartments at the population scale using intact cells. This methodology relies on a multicistronic plasmid design that incorporates a self-cleaving viral 2A peptide to facilitate the production of two separate proteins in transfected cells at a fixed stoichiometry; specifically, a membrane-anchored BRET acceptor (mVenus) and a Luciferase-tagged peripheral lipid-binding probe that serves as the BRET donor. BRET measurements were made at 37°C using a Tristar2 LB 942 Multimode Microplate Reader (Berthold Technologies) with customized emission filters (540/40 nm and 475/20 nm). HEK293-AT1 cells (1×10^5^ cells/well) were seeded in a 200 μL total volume to white-bottom 96 well plates and cultured overnight. Cells were then transfected with 0.25 μg of the specified BRET biosensor using Lipofectamine 2000 (1 μL/well) within OPTI-MEM (40 μL) according to the manufacturer’s protocol; once again with the slight modification of removing the media containing the Lipofectamine-complexed DNA and replacing it with complete culture medium at between 4-6 h post-transfection. Where indicated, additional plasmids, including components of the rapamycin-inducible heterodimerization system, were transfected together with the BRET biosensor at an empirically determined ratio of 1:1:5 for the FRB (0.05 μg), FKBP (0.05 μg), and BRET (0.25 μg) constructs, respectively (0.35 μg/well total). Between 20-24 h post-transfection, the cells were quickly washed before being incubated for 30 minutes in 50 µl of modified Krebs-Ringer buffer (containing 120 mM NaCl, 4.7 mM KCl, 2 mM CaCl_2_, 0.7 mM MgSO_4_, 10 mM glucose, 10 mM HEPES, and adjusted to pH 7.4) at 37°C in a CO_2_-independent incubator. After the pre-incubation period, the cell-permeable luciferase substrate, coelenterazine h (40 µl, final concentration 5 µM), was added and the signal from the mVenus fluorescence and sLuc luminescence were recorded using 485 and 530 nm emission filters over a 4 min baseline BRET measurement (15 s / cycle). Following the baseline recordings, where indicated, the plates were quickly unloaded for the addition of various treatments; which were prepared in a 10 µl volume of the modified Krebs-Ringer solution and added manually. Detection time was always 500 ms for each wavelength and measurements continued for 60 min (15 s / cycle) after the addition of any treatments. All measurements were carried out in triplicate wells and repeated in three independent experiments. From each well, the BRET ratio was calculated by dividing the 530 nm and 485 nm intensities, and normalized to the baseline measurement. The normalized BRET ratio obtained from drug-treated cells was always normalized to vehicle controls. Additionally, to facilitate direct comparisons, were indicated, values obtained from the recruitment of active enzymes were also normalized to control measurements made simultaneously using the corresponding catalytically-inactive variants.

### Protein Expression and Purification

The mutant BcPI-PLC^H82A^ was cloned into a pHIS-GB1 expression vector with an N-terminal His6x-GB1 solubility tag and a TEV cleavage site (Baumlova et al., 2014) The protein was over-expressed in *E.coli* BL21 Star cells and subsequently purified using affinity chromatography on a Ni-NTA resin column (Machery-Nagel). The solubility tag was cleaved with TEV protease and the protein further purified by size exclusion chromatography (Superdex75 GL300/10/300; GE Healthcare) at 4°C in a SEC buffer (20 mM HEPES, 3 mM β-mercaptoethanol, and adjusted pH to 7.0). The purified protein was concentrated to 10 mg/ml and stored at -80°C.

### Crystallization and Crystallographic Analysis

For crystallization, *Bc*PI-PLC^H82A^ (10 mg/ml) was supplemented with 100 mM myo-inositol. The crystals were obtained using the sitting-drop vapor-diffusion method at 18°C by mixing 300 nl of protein-ligand complex with a reservoir solution (Morpheus screen; Molecular dimensions). The diffraction data set was collected at BESSY-II beamline (Mueller et al., 2012). The crystals belonged to the hexagonal P6_3_22 space group and difracted to 2.7Å. Data were proccesed in XDSAPP (Krug et al., 2012) and the structure was solved by molecular replacement using Phaser (McCoy et al., 2007) and previously solved structure of tw PI-PLC (PDB Accession: 1GYM; Heinz et al., 1996) as a search model. Refinement of the structure was performed using Phenix (Adams et al., 2010) or Coot (Debreczeni and Emsley, 2012) to R_work_= 21.62% and R_free_= 24.47%. Protein structures were retrieved using the Protein Data Bank (PDB; Berman et al., 2003) and prepared using the PyMOL Molecular Graphics System Version 2.0 (Schrödinger, LLC).

## Online Supplemental Material

Supplemental Figure S1 compares the overall architecture of the solved *Bc*PI-PLC^H82A^ structure with the existing *myo*-inositol bound *Bc*PI-PLC structure (PDB Accession: 1PTG; Heinz et al., 1995) as well as shows the electron density map surrounding the inositol headgroup in the *Bc*PI-PLC^H82A^ active site. Supplemental Figure S2 compares the steady-state distribution of the *Bc*PI-PLC^H82A^ probe in the COS-7, HEK293-AT1, and HT-1080 cell lineages, with a specific focus on the ER-mitochondria interface. Supplemental Figure S3 provides a schematic explanation of the rationale for using chemical dimerization of the modified FKBP-*Bc*PI-PLC^AA^ to spatially restrict membrane PtdIns hydrolysis and also provides preliminary data directly comparing active site modifications designed to tune the catalysis of the FKBP-*Bc*PI-PLC^AA^ scaffold. Lastly, a catalog of the primers required for PCR-mediated cloning or site-directed mutagenesis are listed in Supplemental Table S5.

## Acknowledgements

This work was supported by the Intramural Research Program (IRP) of the Eunice Kennedy Shriver National Institute of Child Health and Human Development (NICHD). Additional support to J.G.P. was provided by a *Eunice Kennedy Shriver* National Institute of Child Health and Human Development (NICHD) Visiting Fellowship and the Natural Sciences and Engineering Research Council of Canada (NSERC) Banting Postdoctoral Fellowship. A.E. and E.B. were supported from European Regional Development Fund; OP RDE; Project: “Chemical biology for drugging undruggable targets (ChemBioDrug)” (No. CZ.02.1.01/0.0/0.0/16_019/0000729) and by the Academy of Sciences Czech Republic (RVO: 61388963). The authors declare no competing financial interests.

We would like to thank Helmholtz-Zentrum Berlin for the allocation of synchrotron radiation beamtime and the laboratories of Dr. Bianxiao Cui (Stanford University, USA), Dr. Gerry Hammond (University of Pittsburgh, USA), Dr. Takanari Inoue (Johns Hopkins University, USA), Dr. Jennifer Lippincott-Schwartz (HHMI Janelia Research Campus, USA), Dr. Robert Lodge (Institut de Recherches Cliniques de Montreal, Canada), Dr. Harald Stenmark (University of Oslo, Norway), and Dr. Peter Várnai (Semmelweis University Medical School, Hungary) for generously providing DNA constructs. We are also grateful to Dr. Takashi Baba (Akita University, Japan) and Dr. Gergó Gulyás (Semmelweis University Medical School, Hungary) for their assistance with molecular cloning efforts. Confocal imaging was performed at the Microscopy & Imaging Core of the NICHD with the kind assistance of Dr. Vincent Schram and Lynne Holtzclaw. Lastly, with approval from the co-authors, Joshua Pemberton would like to dedicate this study to his Father, the late Dr. S. George Pemberton (University of Alberta, Canada), whose insightful scientific discussions as well as mentorship were invaluable for the completion of this study and the lessons learned will never be forgotten.

## Author Contributions

J.G. Pemberton, Y.J. Kim, D.J. Toth, E. Boura, and T. Balla conceived of the experiments and developed the methods. J.G. Pemberton, Y.J. Kim, N. Sengupta, A. Eisenreichova, E. Boura, and T. Balla performed the experiments. J.G. Pemberton, E. Boura, and T. Balla analyzed the data. T. Balla and E. Boura acquired grant money for this study. J.G. Pemberton wrote the original draft of the manuscript. J.G. Pemberton, Y.J. Kim, E. Boura, and T. Balla reviewed and edited the manuscript.

---

**Supplemental Figure S1.**
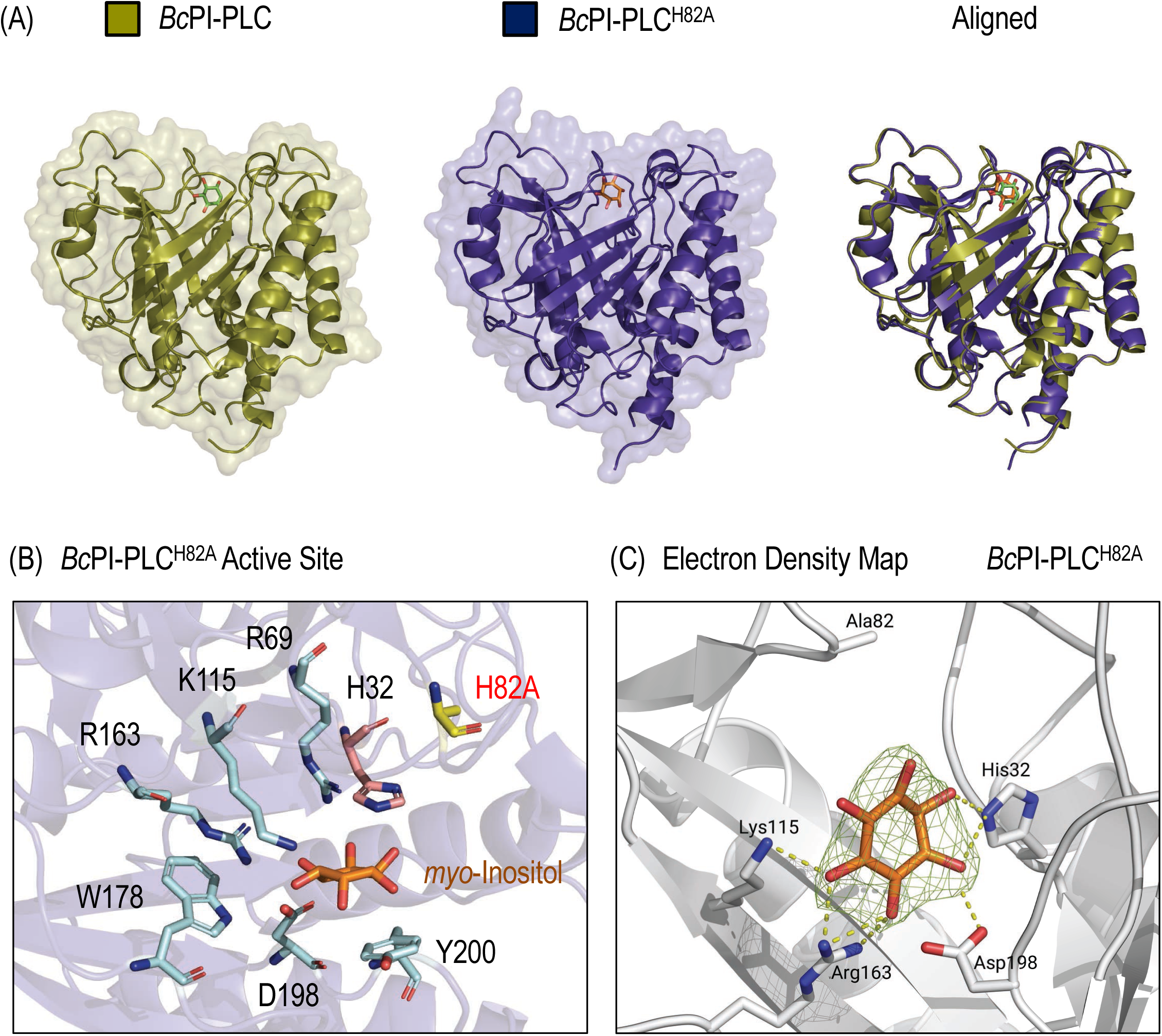
(A) Structural comparison of the *Bc*PI-PLC (gold; PDB Accession: 1PTG; Heinz et al., 1995) and the *Bc*PI-PLC^H82A^ mutant bound to *myo*-inositol. The threaded alignment of the two structures shows no major alterations to the overall protein architecture and only minor changes in the orientation of a few amino acid side chains were observed. (B) Enlarged view of the *Bc*PI-PLC^H82A^ active site, with residues involved in coordinating the *myo*-inositol headgroup highlighted. (C) Electron density map surrounding the *myo*-inositol headgroup within the *Bc*PI-PLC^H82A^ active site; please note the intact coordination of the inositol ring.

**Supplemental Figure S2.**
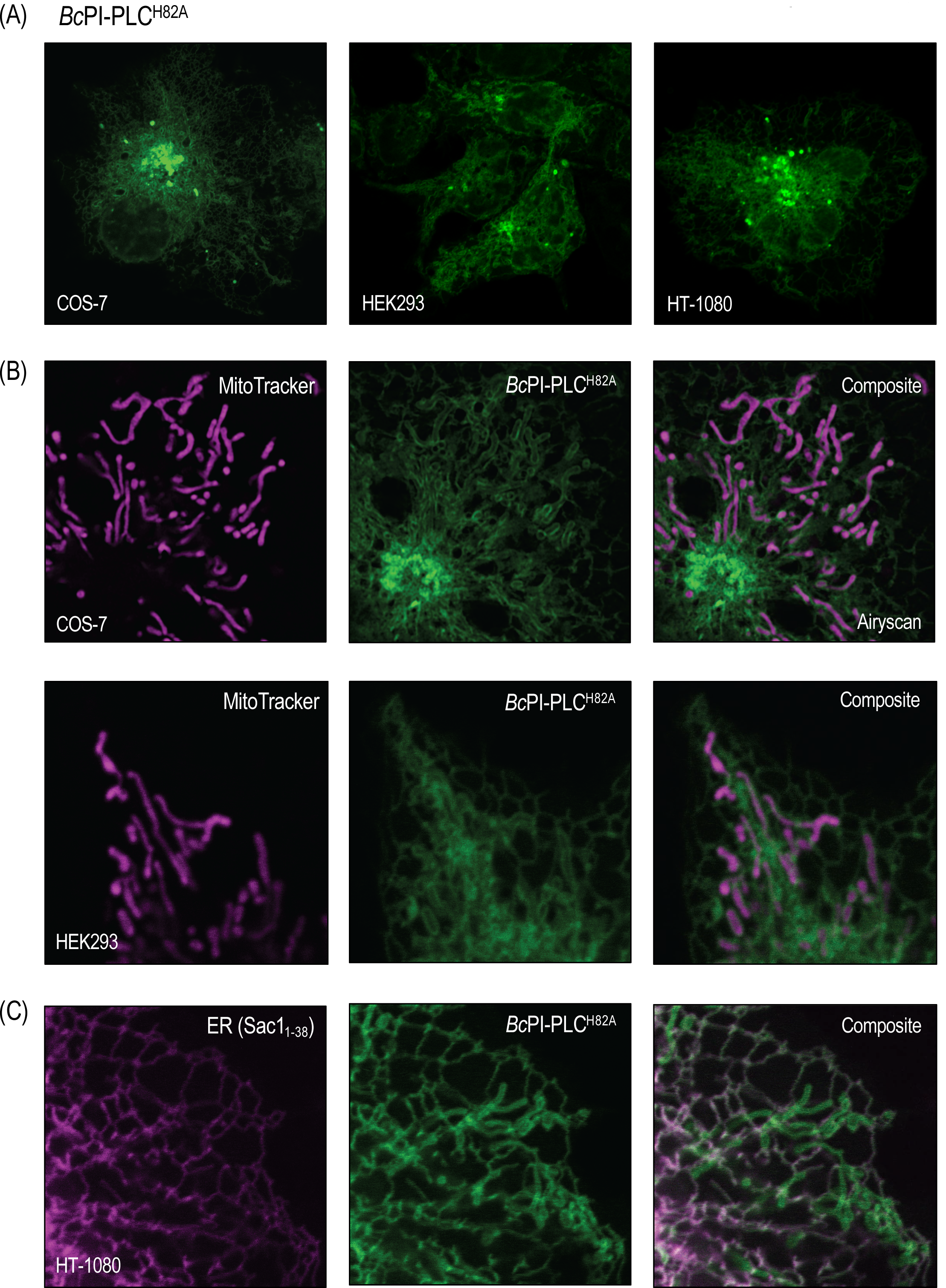
(A) Comparison of the subcellular localization of the EGFP-*Bc*PI-PLC^H82A^ probe in the COS-7, HEK293-AT1, and HT-1080 cell lines. (B) Co-expression of EGFP-*Bc*PI-PLC^H82A^ and the mRFP-tagged signal sequence from the ER-resident protein Sac1 (mRFP-Sac1_1-38_), in HT-1080 cells. (C) Localization of EGFP-*Bc*PI-PLC^H82A^ is shown in HEK293-AT1 (top row) and COS-7 cells (bottom row; super-resolution Airyscan detector) loaded with the lumenal mitochondrial dye, MitoTracker. In general, the steady-state membrane distributions of EGFP-*Bc*PI-PLC^H82A^ were consistent across representative mammalian cell lines and revealed the clear association of the probe with membranes of the ER, mitochondria, and Golgi complex. EGFP-*Bc*PI-PLC^H82A^ also did not localize to the PM or to endosomal compartments in the cell lineages examined.

**Supplemental Figure S3.**
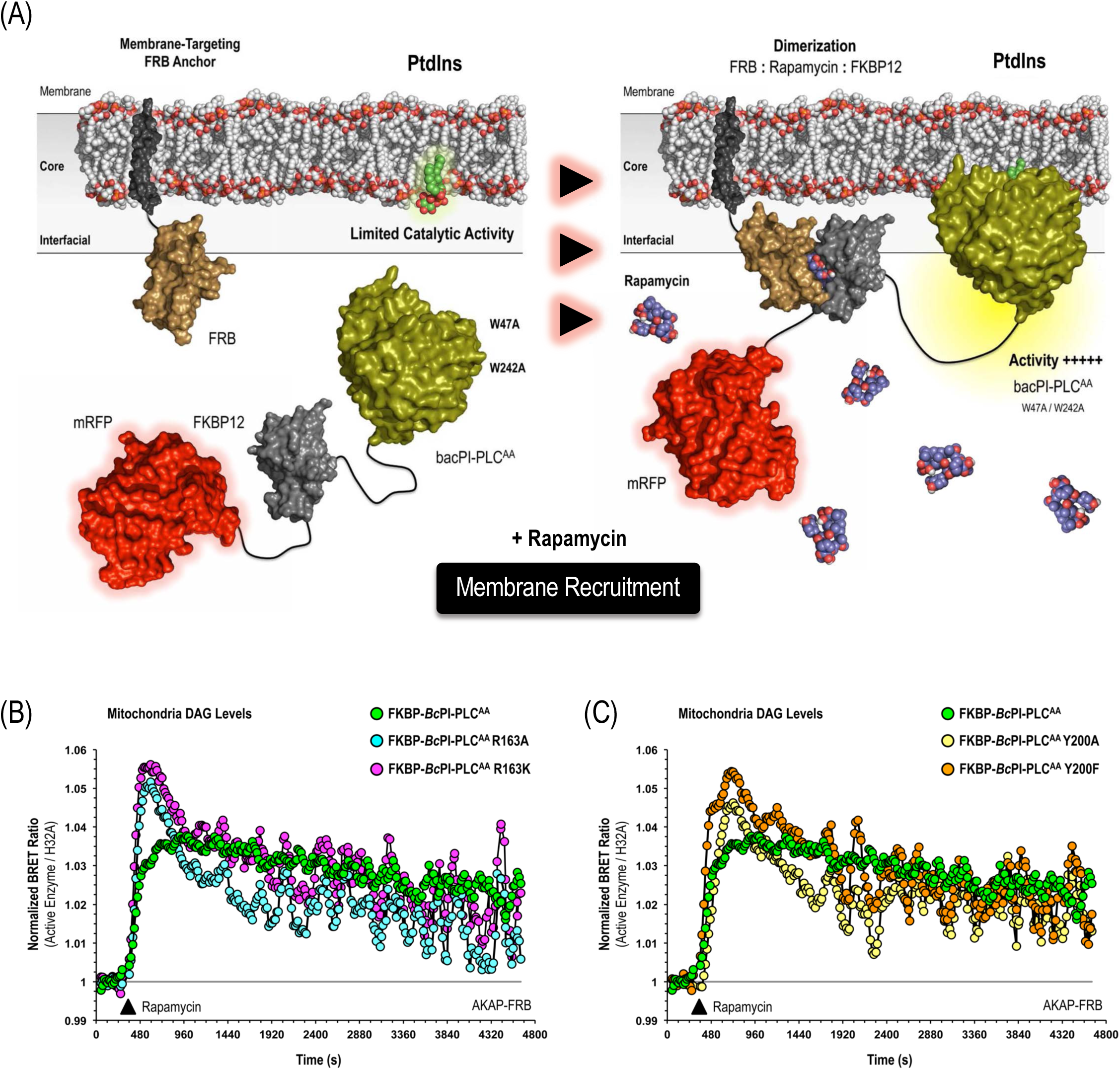
(A) Schematic depicting the use of FRB-tagged membrane anchors to rapidly recruit the FKBP-*Bc*PI-PLC^AA^ scaffold to the surface of specific orgenelles in close proximity to the PtdIns substrate upon acute treatment with rapamycin. Importantly, the rationale mutagenesis of two interfacial tryptophan residues (W47A/W242A; ^AA^) make the resulting FKBP-*Bc*PI-PLC^AA^ enzyme entirely cytosolic, thereby significantly preventing interfacial activation of the enzymatic activity from the cytosol. However, in hopes of further tuning the catalytic activity of the parent FKBP-*Bc*PI-PLC^AA^ scaffold, we screened the activities of mutants with an alanine substitution or conservative mutagenesis of either Arg163 (R163A and R163K; Fig.S3B) or Tyr200 (Y200A and Y200F; Fig.S3C) using the compartment-selective mito-DAG^BRET^ biosensor (AKAP-mVenus-T2A-sLuc-PKDC1_ab_) in combination with a mitochondria-targeted recruiter (AKAP-FRB-ECFP^W66A^). While each of these mutants showed slightly altered kinetics in the DAG-production observed after enzyme recruitment, the rapid initial peak observed for the FKBP-*Bc*PI-PLC^AA^ R163A, R163K, and Y200F variants suggested a minimized consumption of PtdIns by the mutant enzymes before membrane recruitment. To allow for direct comparisons, please note that the pooled response associated with the DAG production stimulated by the parent FKBP-*Bc*PI-PLC^AA^ scaffold is included in both the (B) and (C) panels.

**Table 2:**
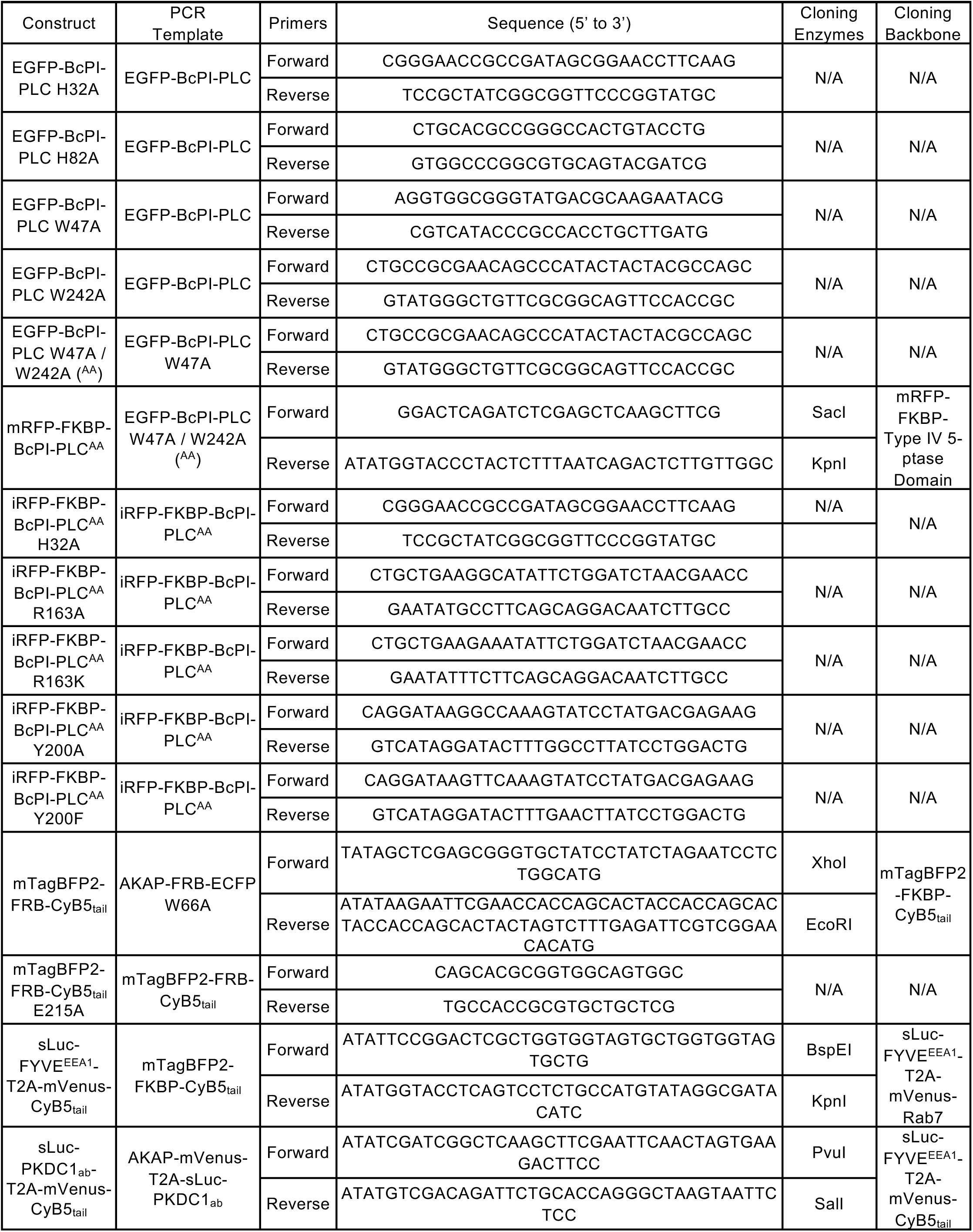
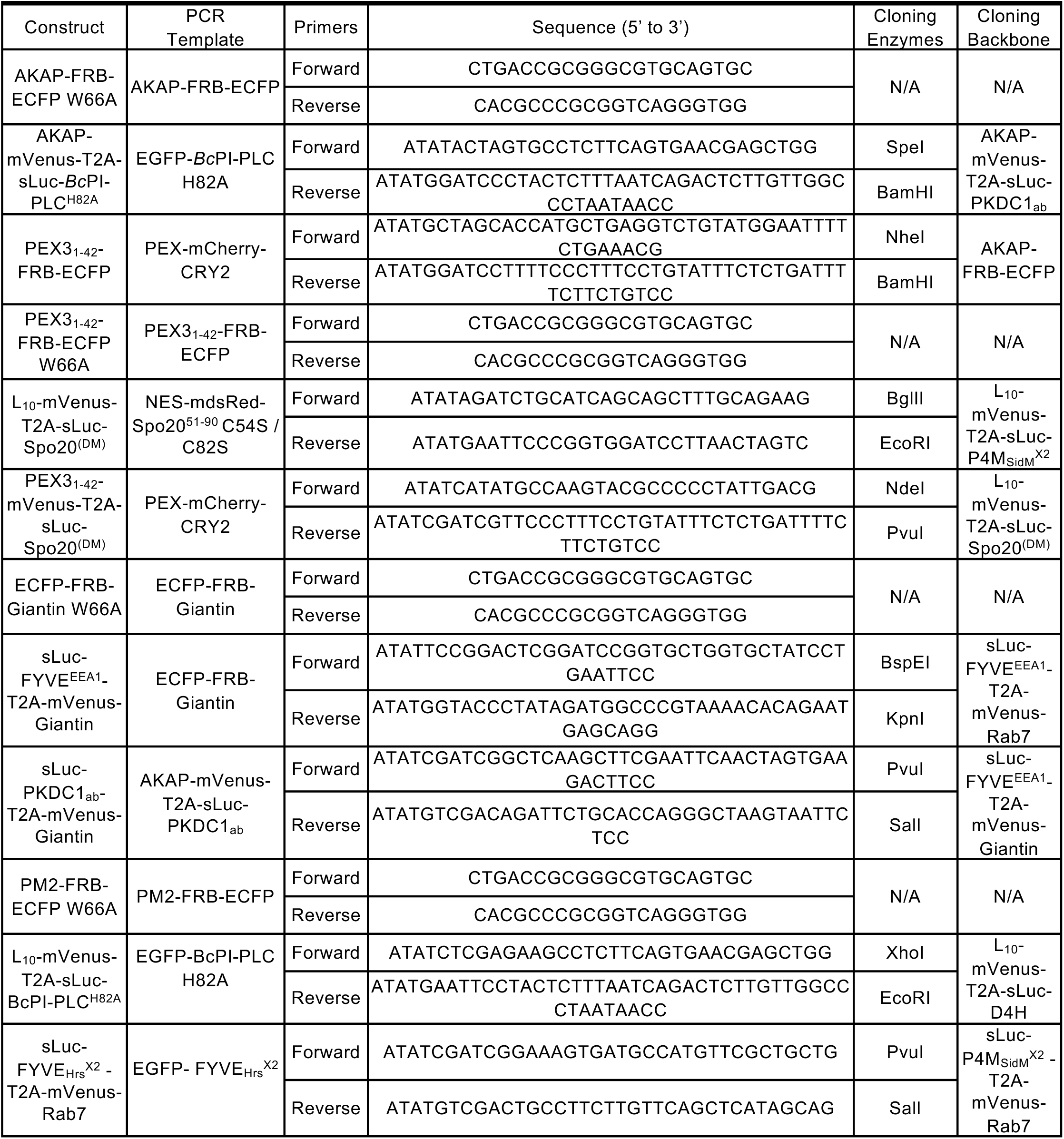
Primers used in this study.

